# When sinks become sources: adaptive colonization in asexuals

**DOI:** 10.1101/433235

**Authors:** Florian Lavigne, Guillaume Martin, Yoann Anciaux, Julien Papaϯx, Lionel Roques

## Abstract

The successful establishment of a population into a new empty habitat outside of its initial niche is a phenomenon akin to evolutionary rescue in the presence of immigration. It underlies a wide range of processes, such as biological invasions by alien organisms, host shifts in pathogens or the emergence of resistance to pesticides or antibiotics from untreated areas.

In this study, we derive an analytically tractable framework to describe the coupled evolutionary and demographic dynamics of asexual populations in a source-sink system. In particular, we analyze the influence of several factors — immigration rate, mutational parameters, and harshness of the stress induced by the change of environment — on the establishment success in the sink (i.e. the formation of a self-sufficient population in the sink), and on the time until establishment. To this aim, we use a classic phenotype-fitness landscape (Fisher’s geometrical model in *n* dimensions) where source and sink habitats determine distinct phenotypic optima. The harshness of stress, in the sink, is determined by the distance between the fitness optimum in the sink and that of the source. The dynamics of the full distribution of fitness and of population size in the sink are analytically predicted under a strong mutation strong immigration limit where the population is always polymorphic.

The resulting eco-evolutionary dynamics depend on mutation and immigration rates in a non straightforward way. Below some mutation rate threshold, establishment always occurs in the sink, following a typical four-phases trajectory of the mean fitness. The waiting time to this establishment is independent of the immigration rate and decreases with the mutation rate. Beyond the mutation rate threshold, lethal mutagenesis impedes establishment and the sink population remains so, albeit with an equilibrium state that depends on the details of the fitness landscape. We use these results to get some insight into possible effects of several management strategies.

## 1 Introduction

Most natural populations are spread over a heterogeneous set of environments, to which local subpopulations may be more or less adapted. When these local populations exchange migrants we can define “source” and “sink” populations. Source populations, where the local genotypes have positive growth rate, are self-sustained and can send migrants to the rest of the system. They may be connected to sink populations, where local genotypes are so maladapted that they have negative growth rates (Pulliam, 1988). A recent review (Furrer and Pasinelli, 2016) showed that empirical examples of sources and sinks exist throughout the whole animal kingdom. In the absence of any plastic or evolutionary change, source-sink systems are stable, with the sources being close to their carrying capacity and the sinks being only maintained by incoming maladapted migrants from source environments. In the literature, different source-sink systems have been categorized by their pattern of immigration and emigration (for more detail on these different categories see Fig. 1 in Sokurenko et al. (2006) and Table 1 in Loreau et al. (2013)). One particular system, defined as “black-hole sink” (Gomulkiewicz et al., 1999), corresponds to a demographic dead-end, from which emigration is negligible. These black-hole sinks, and their demographic and evolutionary dynamics, are the canonical model for studying the invasion of a new environment, outside of the initial species “niche”, and thus initially almost empty (Holt et al., 2003, 2004). In this article, we will only consider black-hole sinks: for compactness, we hereafter simply use the term ‘sink’, when in fact referring to a black-hole sink population. The demographic and evolutionary process leading, or not, to the invasion of a sink is akin to evolutionary rescue in the presence of immigration. It underlies a wide range of biological processes: invasion of new habitats by alien organisms (Colautti et al., 2017), host shifts in pathogens or the emergence of resistance to pesticides or antibiotics, within treated areas or patients (discussed e.g. in Jansen et al. (2011) and Sokurenko et al. (2006)). The issues under study in these situations are the likelihood and timescale of successful invasions (or establishment) of sinks from neighboring source populations. “Establishment” in a sink is generally considered successful when the population is self-sustaining in this new environment, even if immigration was to stop (e.g., Blackburn et al., 2011, for a definition of this concept in the framework of biological invasions).

**Table 1:**
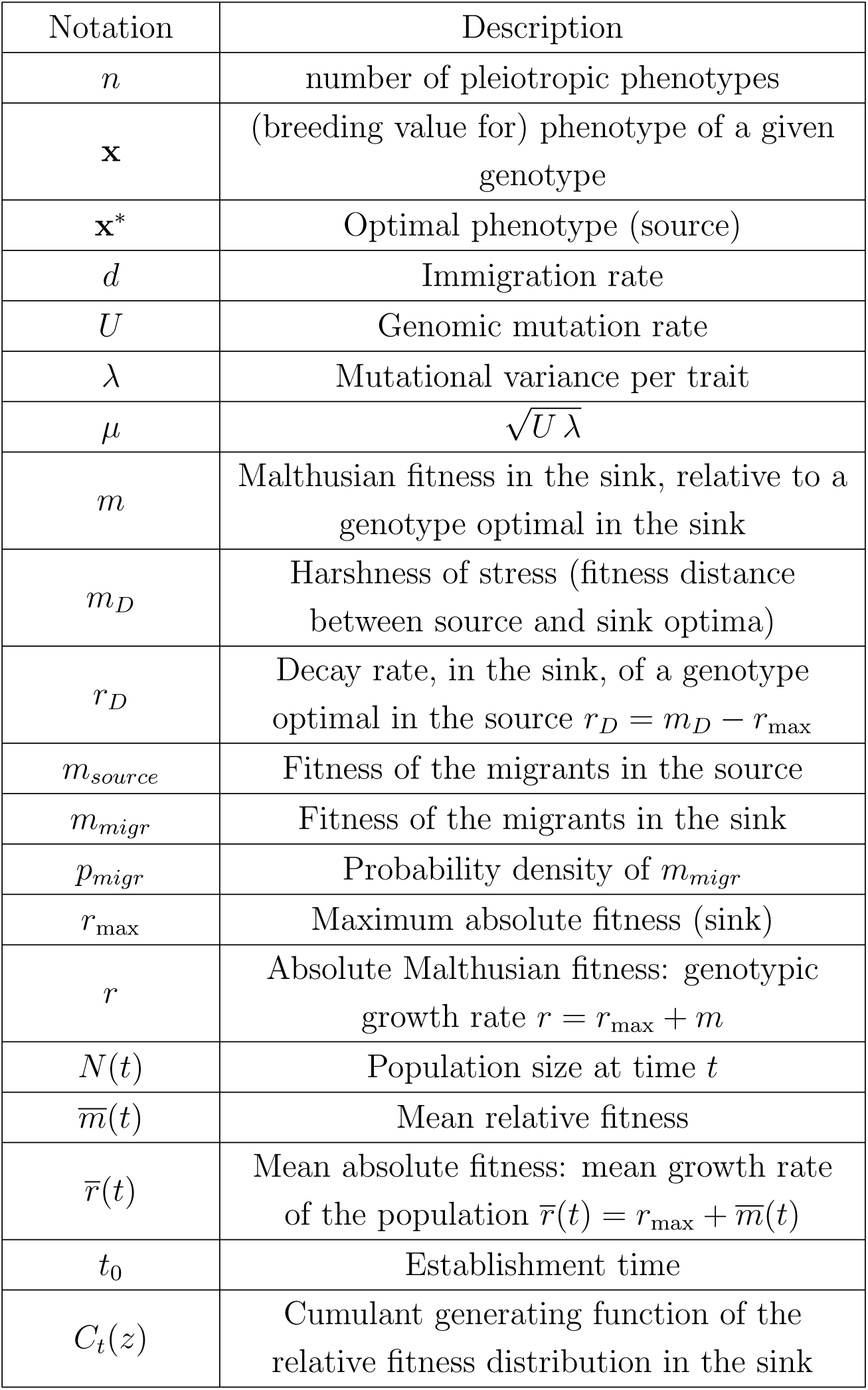
Main notations.

A rich theoretical literature has considered the effects of demography and/or evolution in populations facing a heterogeneous environment connected by migration, both in sexuals (e.g., Kirkpatrick and Barton, 1997) and asexuals (e.g., Débarre et al., 2013). The source-sink model is a sub-case of this general problem, that has received particular attention (for a review, see Holt et al., 2005): below, we quickly summarize the relevance and key properties of source-sink models. The asymmetric migration (from source to sink alone), characteristic of black-hole sinks, provides a key simplification, while remaining fairly realistic over the early phase of invasion, where success or failure is decided. For the same reason, some models further ignore density-dependent effects in the sink, although both high (logistic growth) and/or low (Allee effect) densities could further impact the results, when relevant (discussed in Holt, 2009).

Some source-sink models (e.g., Drury et al., 2007; Garnier et al., 2012), focus on detailed demographic dynamics, in the absence of any evolutionary forces. Evolutionary forces (selection, mutation, migration, drift and possibly recombination/segregation) can greatly alter the outcome. These forces may yield both local adaptation or maladaptation, favoring or hindering (respectively) the ultimate invasion of the sink (“adaptive colonization”, Gomulkiewicz et al., 2010), however harsh. In this context, mutation and migration are double edged swords, both increasing the local variance available for selection but generating mutation and migration loads, due to the adverse effects of deleterious mutations and maladapted migrant inflow (resp.). For a review of the ambivalent effects of mutation and migration see e.g., (Lenormand, 2002) and (Débarre et al., 2013). Disentangling the complex interplay of these forces with demographic dynamics is challenging, and modelling approaches have used various ecological simplifications: e.g. no age or stage structure, constant stress, constant migration rate.

The associated evolutionary processes are also simplified. As for evolutionary rescue models (discussed in Alexander et al., 2014), evolutionary source-sink models may be divided into two classes, based on the presence or absence of context-dependence in the genotype-fitness map they assume (Gomulkiewicz et al., 2010). In context-independent models, fitness in the sink is additively determined by a single or a set of freely recombining loci, and adaptation occurs by directional selection on fitness itself (Gomulkiewicz et al., 2010; Barton and Etheridge, 2017). In context-dependent models, which arguably forms the vast majority of source-sink models, fitness is assumed to be a concave function (typically quadratic or Gaussian) of an underlying phenotype, with the source and sink environments corresponding to alternative optima for this phenotype (e.g., Holt et al., 2003, 2004). Such nonlinear phenotype-fitness maps, with environment dependent optima, generate context-dependent interactions for fitness (epistasis and genotype x environment or “G x E” interactions): the effect of a given allele depends on the genetic and environmental background in which it is found. These models reproduce observed empirical patterns of mutation fitness effects across backgrounds (Martin et al., 2007; MacLean et al., 2010; Trindade et al., 2012), reviewed in (Tenaillon, 2014). However, their analysis is more involved. Most analytical treatments have thus relied on stationarity assumptions: e.g. describing the ultimate (mutation-selection-migration) equilibrium in asexuals (Débarre et al., 2013), or assuming a constant genetic variance and Gaussian distribution for the underlying trait in sexuals (e.g., Gomulkiewicz et al., 1999; Holt et al., 2004). While numerical explorations (by individual-based simulations) often relax these stationarity assumptions, they are necessarily bound to study a limited set of parameter value combinations.

In this paper, we explore a complementary scenario: a source-sink system, out of equilibrium, in an asexual population. The focus on asexuals is intended to better capture pathogenic microorganisms or microbial evolution experiments. We ignore density-dependence by assuming that it is negligible (no Allee effects) before and during the critical early phase of the sink invasion (far below the population reaches the carrying capacity). Considering asexuals and density-independent populations implies that several complex effects of migration (both genetic and demographic) can be ignored. Because migrants do not hybridize/recombine with locally adapted genotypes or use up limiting resources, the maladaptive effects of migration are limited. Migration meltdown and gene swamping (see Lenormand, 2002) are thus expected to be absent. This simplification allows to analytically track out-of-equilibrium dynamics, in a context-dependent model (with epistasis and G x E), without requiring stable variance or Gaussian and moment-closure approximations for the phenotypic distribution.

More precisely, we study the transient dynamics of a sink under constant immigration from a source population at mutation-selection balance and a sink initially empty (invasion process). We use the classic quadratic phenotype-fitness map with an isotropic version of Fisher’s geometrical model (FGM) with mutation pleiotropically affecting *n* phenotypic traits. To make analytical progress, we use a deterministic approximation (as in Martin and Roques, 2016) that neglects stochastic aspects of migration, mutation and genetic drift, but tracks the full distribution of fitness and phenotypes. Under a weak selection strong mutation (WSSM) regime, when mutation rates are large compared to mutation effects, we further obtain an analytically tractable coupled partial-ordinary differential equation (PDE-ODE) model describing the evolutionary and demographic dynamics in the sink. This framework allows us to derive analytic formulae for the demographic dynamics and the distribution of fitness, at all times, which we test by exact stochastic simulations. We investigate the effect of demographic and evolutionary parameters on the establishment success, on the establishment time, and on the equilibrium mean fitness in the sink. In particular, we focus on the effects of the immigration rate, the harshness of stress (distance between source and sink optima), and mutational parameters (rate, phenotypic effects and dimension *n*).

## 2 Methods

Throughout this paper, we follow the dynamics of the fitness distribution of the individuals in the sink environment, under the joint action of mutation, selection and immigration from the source. The latter remains stable at mutation-selection balance, as migration is asymmetric in this black-hole sink. We consider an asexual population evolving in continuous time. Consistently, we focus on Malthusian fitness *m* (hereafter ‘fitness’): the expected growth rate (over stochastic demographic events) of a given genotypic class, per arbitrary time units. Absolute Malthusian fitnesses *r* are therefore (expected) growth rates, and without loss of generality, *m* is measured relative to that of the phenotype optimal in the sink, with growth rate *r*_max_. We thus have *m* = *r − r*_max_, and the mean absolute fitness 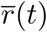 and mean relative fitness 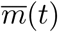, at time *t*, satisfy:

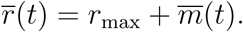

We use a *deterministic approximation* which neglects variations among replicate populations. Under this approximation, 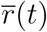 (respectively 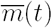), the mean absolute (resp. relative) fitness within each population can be equated to their expected values (across stochastic events). In general, the bar ¯ denotes averages taken over the sink population. The main notations are summarized in Table 1.

### 2.1 Demographic model and establishment time *t*_0_

In our simple scenario without density-dependence, evolutionary and demographic dynamics are entirely coupled by the mean absolute Malthusian fitness (mean growth rate). We consider a sink population with mean growth rate 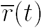 at time *t*, receiving on average *d* individuals per unit time by immigration. Under the deterministic approximation, the population size dynamics in the sink environment are therefore given by:

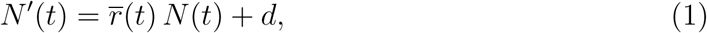

with *N′* (*t*) the derivative of *N* with respect to *t* at time *t*.

In the absence of adaptation, 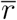 is constant, leading to an equilibrium population size 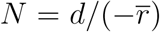 when 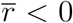, as mentioned in the Introduction. When genetic adaptation is taken into account, we need further assumptions to describe the dynamics of 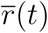 in the sink.

We always assume that the new environment is initially empty (*N* (0) = 0) and that the individuals from the source are, on average, maladapted in the sink 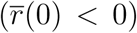. Following a classic definition (Blackburn et al., 2011), we define the establishment time *t*_0_ as the first time when the growth rate of the sink becomes positive in the absence of immigration:

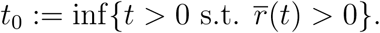

This means that, from time *t*_0_, the sink population is self-sustaining in the absence of immigration and further adaptation. By definition (assuming that 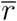 is continuous), *t*_0_ satisfies 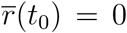. Depending on the behavior of *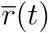, t*_0_ may therefore be finite (successful establishment) or infinite (establishment failure).

### 2.2 Fisher’s geometric model

We use Fisher’s geometric model (FGM) to describe the relationships between genotypes, phenotypes and fitnesses in each environment. This phenotype-fitness landscape model has the advantage of yielding realistic distributions of mutation effects on fitnesses (Trindade et al., 2012; Hietpas et al., 2013; Tenaillon, 2014) and of generating a coupling between stress levels, the distribution of fitnesses among migrants from the source and that among *de novo* random mutants arising in the sink (Anciaux et al., 2018).

#### Phenotype-fitness relationships in the two environments

The FGM assumes that each genotype is characterized by a given breeding value for phenotype at *n* traits (hereafter simply denoted ‘phenotype’), namely a vector **x** *∈* ℝ ^*n*^. Each environment (the source and the sink) is characterized by a distinct phenotypic optimum. The distance between these optima determines the stress induced by a change of the environment. An optimal phenotype in the sink has maximal absolute fitness *r*_max_ (relative fitness *m* = 0) and sets the origin of phenotype space (**x** = 0). Fitness decreases away from this optimum. Following the classic version of the FGM, Malthusian fitness is a quadratic function of the breeding value *r*(**x**) = *r*_max_ −‖**x**‖^2^*/*2 and *m*(**x**) = −‖**x**‖^2^*/*2.

In the source, due to a different phenotype optimum **x***\* ∈* ℝ ^*n*^, the relative fitness is *m\**(**x**) = −‖**x**−**x***\**‖^2^*/*2. As the population size is kept constant in the source (see below), only relative fitness matters in this environment. The harshness of stress *m*_*D*_ *>* 0 is the fitness distance between source and sink optima:

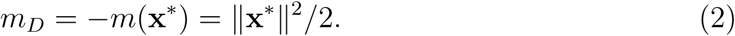

The decay rate, in the sink, of a population composed of individuals with the optimal phenotype from the source, is thus *r*_*D*_ = *m*_*D*_ − *r*_max_.

#### Mutations

In the two environments, mutations occur at rate *U* and create independent and identically distributed (iid) random variations **dx** around the phenotype of the parent, for each trait. We assume here a standard Gaussian distribution of the mutation phenotypic effects (Kimura, 1965; Lande, 1980): **dx** *∼ 𝒩* (0, *λI*_*n*_), where *λ* is the mutational variance at each trait, and *I*_*n*_ is the identity matrix in *n* dimensions. These assumptions induce a distribution of the mutation effects on fitness, given the relative fitness *m*_*p*_ *≤* 0 of the parent. This distribution has stochastic representation (Martin, 2014) 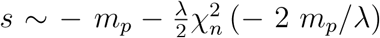, where 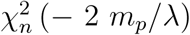 denotes the noncentral chi-square distribution with *n* degrees of freedom and noncentrality −2 *m*_*p*_*/λ*. This distribution is detailed elsewhere (reviewed in Tenaillon, 2014), its mean is 𝔼[*s*] = −*n λ/*2. Alternatively, it can be characterized by its moment generating function:

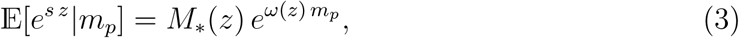

with

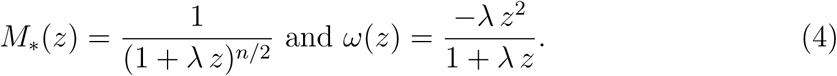

#### Migration events

Migration sends randomly sampled individuals from the source into the sink, at rate *d >* 0 per unit time. Their relative fitness in the sink is *m*_*migr*_(**x**) = −‖**x**‖^2^*/*2, with **x** randomly sampled from the source’s standing phenotype distribution.

### 2.3 Fitness distribution of the migrants

We assume that the distribution of phenotypes in the source is at mutation-selection balance. The resulting equilibrium distribution of phenotypes yields an equilibrium fitness distribution in the source. Under a weak selection strong mutation (WSSM) regime, a simple expression for this equilibrium fitness distribution is (Martin and Roques, 2016, equation (10)): *m*_*source*_ *∼ −*Γ(*n/*2, *µ*), with 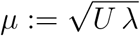, where Γ(*a, b*) denotes a gamma deviate with shape *a* and scale *b*. This WSSM regime can be quantitatively defined by the inequality *U > U*_*c*_ := *n*^2^ *λ/*4 (Martin and Roques, 2016, Appendix E).

To understand the dynamics of the fitness distribution in the sink, we need to compute the distribution of the relative fitness of the migrants *m*_*migr*_ when they arrive into the sink. In our case, a handy way to describe this distribution is to compute its moment generating function: 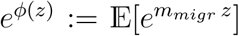, for any *z ≥* 0. Computations in Appendix A show that for any *z ≥* 0:

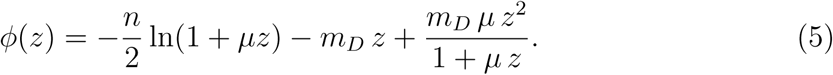

The corresponding distribution of *m*_*migr*_ (see Appendix A) is:

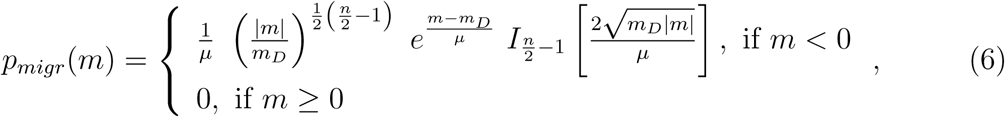

where *I*_*v*_ is the modified Bessel function of the first kind. The accuracy of this formula is illustrated in Fig. 1. We observe that the mean absolute fitness of the migrants, which coincides with 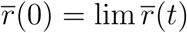 as *t* → 0, is given by

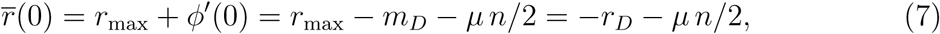

with *ϕ* defined by (5). This initial growth rate is negative and corresponds to the decay rate (*r*_*D*_) of the mean phenotype from the source (which is optimal there) plus a variance load (*µ n/*2) due to the equilibrium variation around this mean.

The assumption that the individuals from the source are initially decaying 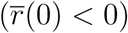 can therefore be expressed by the inequality *r*_max_ − *µ n/*2 *< m*_*D*_.

### 2.4 Trajectories of fitness in the sink: a PDE approach

At time *t*, the population in the sink consists of the phenotypes {**x**_*i*_(*t*)}_*i*=1,*…,N*(*t*)_ (with *N* (*t*) *∈* ℕ), with the corresponding values of relative fitnesses {*m*_*i*_(*t*)}_*i*=1,*…,N*(*t*)_. In the absence of demography and immigration, the dynamics of the fitness distribution is traditionally investigated by a moment closure approximation (Burger, 1991; Gerrish and Sniegowski, 2012): the variations of the moment of order *k* depend on the moments of order larger than (*k* + 1) through a linear ordinary differential equation, and the resulting system is solved by assuming that the moments vanish for *k* larger than some value. A way around this issue is the use of cumulant generating functions (CGFs), which handle all moments in a single function. In a relatively wide class of evolutionary models of mutation and selection, the CGF of the fitness distribution satisfies a partial differential equation (PDE) that can be solved without requiring a moment closure approximation (Martin and Roques, 2016, Appendix B). We follow this approach here. The empirical CGF of the relative fitness in a population of *N* (*t*) individuals with fitnesses *m*_1_(*t*), *…, m*_*N*(*t*)_(*t*) is defined by

**Figure 1:**
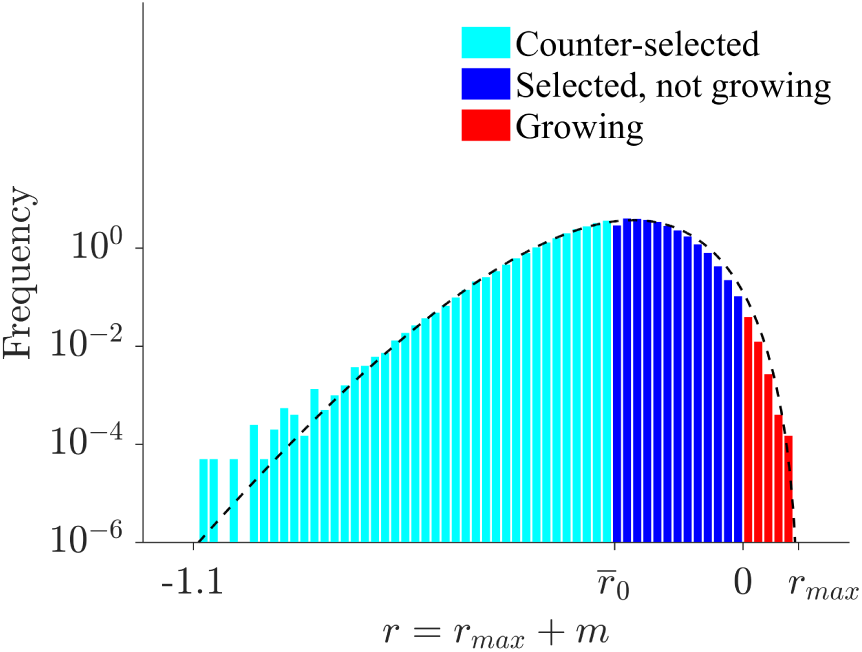
Distribution of absolute fitness of the migrants in the sink. The dashed line corresponds to the theoretical expected values of this distribution *p*_*migr*_(*· - r*_max_) given by formula (6). The histogram corresponds to the distribution of migrants obtained in exact stochastic simulations after reaching the mutation-selection balance in the source (see Section 2.5). When the sink is empty, individuals are ‘counter-selected’ if their fitness is below the mean fitness 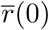 given by (7), ‘selected’ if their fitness is above 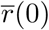, and ‘growing’ if their fitness is positive. The parameter values are *r*_max_ = 0.1, *U* = 0.1, *m*_*D*_ = 0.3, *λ* = 1*/*300, *n* = 6 and *N* = 10^6^.

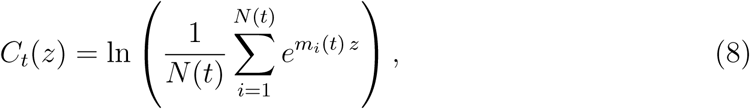

for all *z ≥* 0. The mean fitness and the variance in fitness in the sink can readily be derived from derivatives, with respect to *z*, of the CGF, taken at *z* = 0: 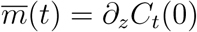 (and 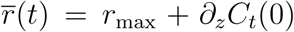), and *V* (*t*) = *∂*_*zz*_*C*_*t*_(0) (the variance in fitness). In the absence of demography and immigration, and under a weak selection strong mutation (WSSM) regime, (Martin and Roques, 2016, Appendix A) derived a deterministic nonlocal PDE for the dynamics of *C*_*t*_. We extend this approach to take into account immigration effects and varying population sizes. This leads to the following PDE (derived in Appendix B):

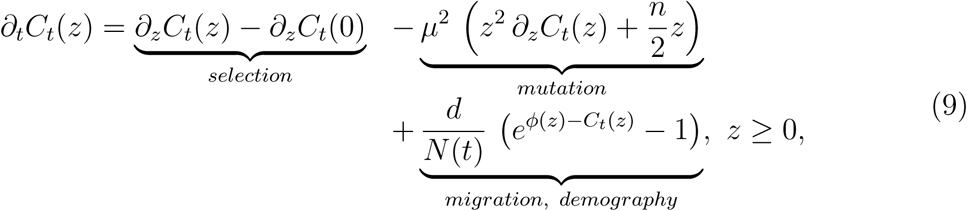

where we recall that 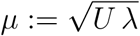. The immigration term depends on *ϕ*(*z*), which is given by (5), and on *N* (*t*), which satisfies the ODE (1), i.e. *N′* (*t*) = (*∂*_*z*_*C*_*t*_(0)+ *r*_max_) *N* (*t*)+ *d.* This leads to a well-posed coupled system (1) & (9) which can be solved explicitly, as shown in Appendix C.

The selection term in eq. (9) stems from the increase in frequency of each lineage proportionally to its Malthusian fitness (frequency-independent selection). The second term is the WSSM approximation (*U > U*_*c*_) to a more complex term (Martin and Roques, 2016, Appendix A) describing the effect of mutation: it depends on the current background distribution (on *C*_*t*_(*z*)) because of the fitness epistasis inherent in the FGM. The last term describes the effect of the inflow of migrants on lineage frequencies. It tends to equate *C*_*t*_(*z*) with *ϕ*(*z*), the CGF of fitnesses among migrants, proportionally to *d/N* (*t*), the dilution factor of migrants into the current sink population.

### 2.5 Individual-based stochastic simulations

To check the validity of our approach, we used as a benchmark an individual-based, discrete time model of genetic drift, selection, mutation, reproduction and migration with non-overlapping generations.

#### Source population

A standard Wright-Fisher model with constant population size was used to compute the equilibrium distribution of phenotypes in the source. Our computations were carried out with *N \** = 10^6^ individuals in the source. Each individual *i* = 1, *…, N \** has phenotype **x**_*i*_ *∈*ℝ^*n*^ and relative Malthusian fitness *m*_*i*_ = −‖**x**_*i*_ − **x***\**‖^2^*/*2, with corresponding Darwinian fitness 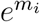 (discrete time counterpart of the Malthusian fitness). At each generation, *N \** individuals are sampled with replacement proportionally to their Darwinian fitness. Mutations are simulated by randomly drawing, every generation and for each individual, a Poisson number of mutations, with rate *U*. Mutation acts additively on phenotype, with individual effects *d***x** drawn into an isotropic multivariate Gaussian distribution with variance *λ* per trait (see Section 2.2). Simulations were started with a homogeneous population (**x**_*i*_ = **x***\** for all *i* at initial time) and ran for 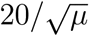 generations (the predicted time taken to reach a proportion *q* of the final equilibrium mean fitness is 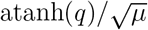, see Appendix E, Section “Characteristic time” in Martin and Roques (2016); with atanh(*q*) = 20, one can consider that the equilibrium has been reached). An example of the distribution of absolute fitness in the resulting (equilibrium) source population, after migrating into the sink (distribution of *r*_max_ −‖**x**_*i*_‖^2^*/*2) is presented in Fig. 1.

#### Sink population

We started with *N* (0) = 0 individuals in the sink. Then, the process to go from generation *t* to generation (*t* + 1) is divided into three steps: (i) migration: a Poisson number of migrants, with rate *d*, was randomly sampled from the equilibrium source population, and added to the population in the sink; (ii) reproduction, selection and drift: each individual produced a Poisson number of offspring with rate exp(*r*_*i*_) = exp(*r*_max_ + *m*_*i*_) (absolute Darwinian fitness in the sink); (iii) mutation followed the same process as in the source population. The stopping criterion was reached when *N* (*t*) *>* 1.5 *·* 10^6^ individuals or *t >* 5 *·* 10^3^ to limit computation times.

All the Matlab^®^ codes to generate individual-based simulations are provided in Supplementary File 1.

## 3 Results

### 3.1 Trajectories of mean fitness

#### Dynamics of 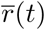 and *N* (*t*)

The system (1) & (9) leads to an expression for the mean absolute fitness (Appendix C):

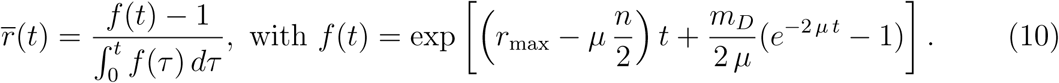

It also leads to an expression for the population size thanks to 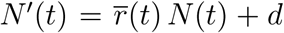. (see eq. (16) in Appendix C).

The good accuracy of eq. (10) is illustrated in Figs. 2-4, by comparing it with the results of individual-based stochastic simulations, under the WSSM assumption (*U > U*_*c*_ := *n*^2^ *λ/*4). Both the individual-based simulations and the analytic expressions show that sink invasion tends to follow four different phases, which are all the more pronounced as the harshness of stress *m*_*D*_ increases. *Phase 1:* During the first generations, the mean fitness slightly increases; *Phase 2:* The mean fitness remains stable. *Phase 3:* Rapid increase in mean fitness. *Phase 4:* The mean fitness stabilizes at some asymptotic value. In the case of establishment failure (Fig. 4), the adaptation process remains in Phase 2.

In all cases, formula (7) gives an accurate prediction of the mean fitness of the migrants, as shown by the agreement between theoretical and simulated values of 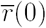. Other trajectories, outside of the WSSM regime (*U < U*_*c*_) are presented in Appendix D (and discussed in Section 3.3).

#### Phenotypic dynamics over the different phases of invasion

Obviously the dichotomy into four phases could be deemed somewhat arbitrary, and it is clearly less marked with milder stress (top panels of Fig. 2). However, it does convey the qualitative chronology of the whole process in all cases. This can be further understood by exploring the dynamics of the phenotypic distribution over time: a typical example for a single simulation is given in Fig. 3, at four times corresponding to each of the four phases. We show here the phenotypic distribution along the one meaningful dimension, that for which the optimum is shifted between source and sink (the optimum in the sink is 0, and the optimum in the source 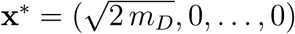). The corresponding trajectories of fitness and population size are available in Appendix E (Fig. 9). A video file of the phenotype distribution is also available as Supplementary File 2.

During Phases 1 and 2, the phenotypic distribution is fairly stable and slightly shifted from the source distribution towards the sink optimum. The short Phase 1 merely witnesses an increase in population size from zero to the semi-stable Phase 2. We suggest that this semi-stable state approximately corresponds to a macroscopic “equilibrium” between migration and selection on the bulk of phenotypes. Here, we conjecture a negligible impact of mutation on this bulk because simulations in the absence of mutation in the sink yield a very similar phenotypic distribution during Phase 2 (Appendix J, Fig. 12). However, over the course of Phase 2, a second mode slowly appears closer to the sink optimum, due to the invasion of rare, better adapted, phenotypes (generated by the combined effects of rare adapted migrants and *de novo* mutation in the sink). When this second mode becomes significant in frequency, Phase 3 starts with a rapid increase of the second mode (and of mean fitness), because phenotypic and fitness variance are then maximized. The last Phase 4 corresponds to the new equilibrium dominated by a mutation selection balance around the sink optimum. In the present model without density limitations, migration becomes ultimately negligible as the sink population explodes, and its phenotypic distribution ultimately reaches exactly a new mutation-selection balance.

**Figure 2:**
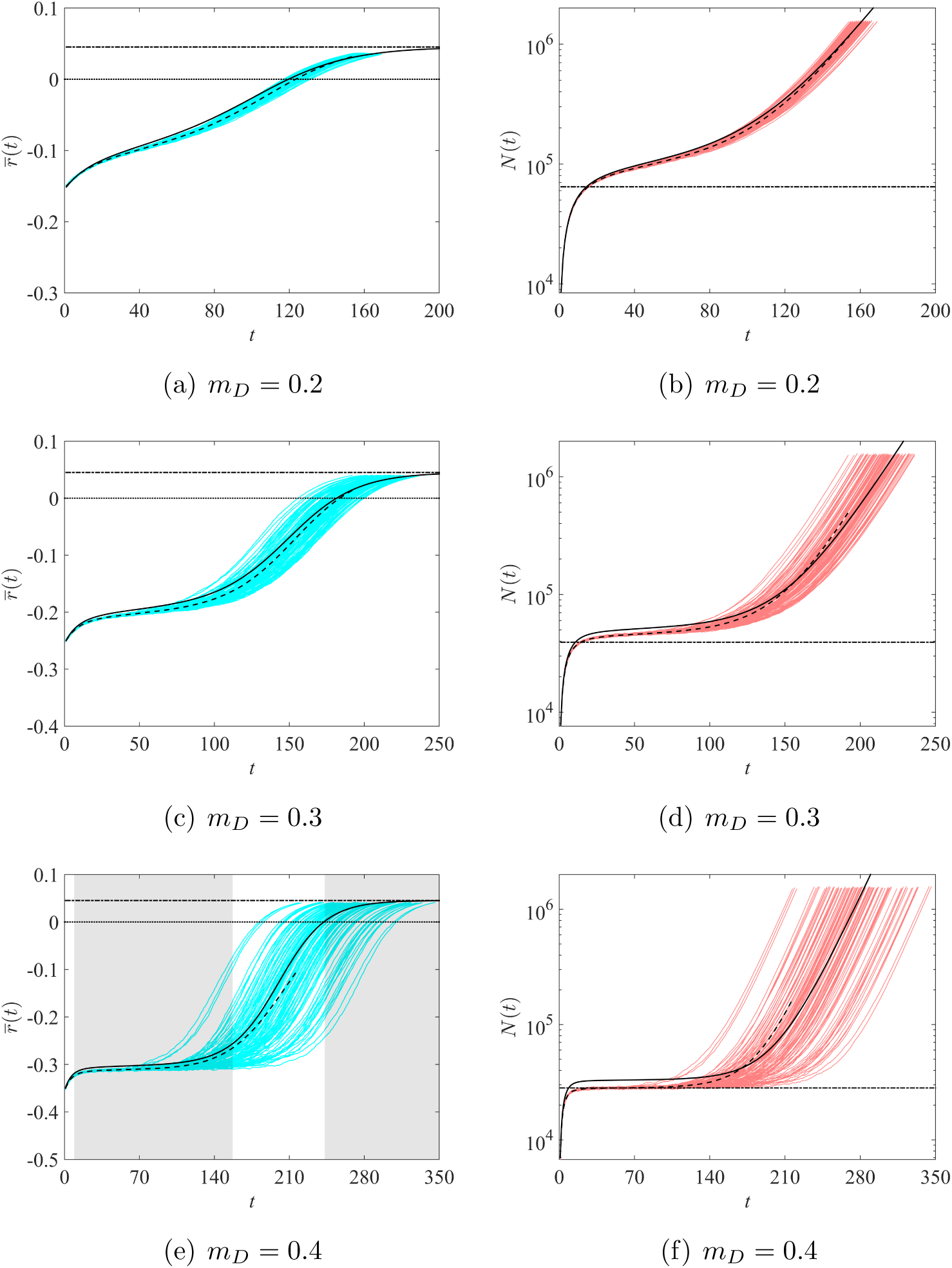
Trajectories of mean fitnesses and population sizes in a WSSM regime, depending on the harshness of stress. Solid lines: analytical predictions given by formulae (1) and (10) vs 100 trajectories obtained by individual-based simulations (blue curves for 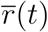 and red curves for *N* (*t*); dashed lines: mean values averaged over the 100 populations). Horizontal dashed-dotted lines: theoretical value of 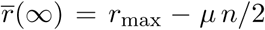 (left panels) and equilibrium population size 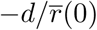 in the absence of adaptation (right panels). The four phases of invasion (Phases 1-4, see main text) are illustrated by distinct shaded areas on panel (e). The parameter values are *U* = 0.1 (thus, *U > U*_*c*_ = 0.03, which is consistent with the WSSM regime), *r*_max_ = 0.1, *λ* = 1*/*300, *n* = 6 and *d* = 10^4^. Due to the stopping criterion *N* (*t*) = 1.5 *·* 10^6^ was reached, the mean values could not be computed over the full time span.

**Figure 3:**
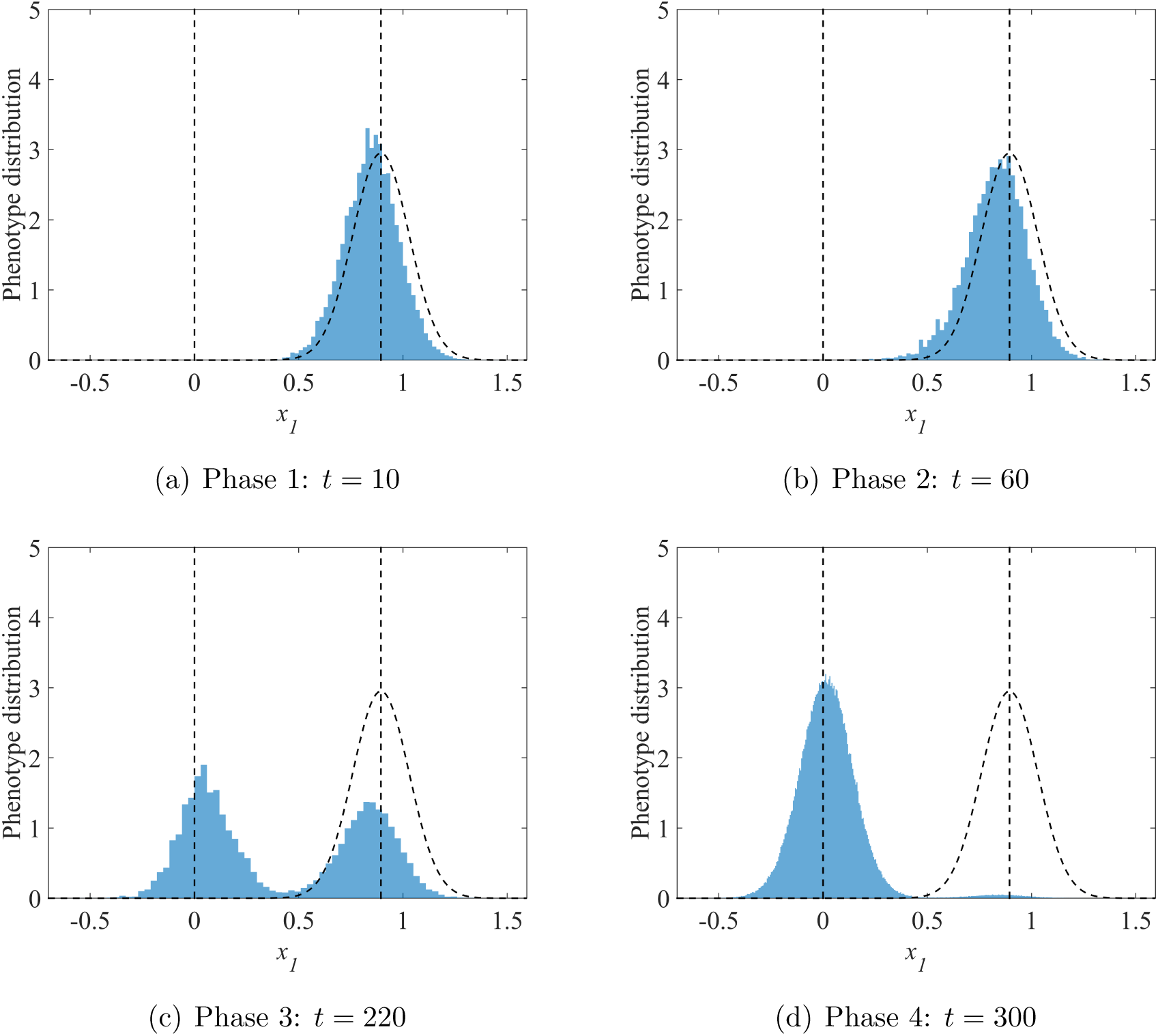
Phenotype distribution in the sink, along the direction *x*_1_. The vertical dotted lines correspond to the sink (*x*_1_ = 0) and source 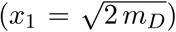 optima. The black dotted curve corresponds to the theoretical distribution of migrant’s phenotypes in the sink (Gaussian distribution, centered at 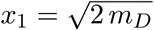, and with variance 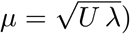). In all cases, the parameter values are *m*_*D*_ = 0.4, *U* = 0.1, *r*_max_ = 0.1, *λ* = 1*/*300, *n* = 6 and *d* = 10^4^.

#### Effect of the immigration rate

Unexpectedly, the value of 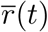 in formula (10) does not depend on the immigration rate *d*. Thus, only the population size dynamics are influenced by the immigration rate, but not the evolutionary dynamics. To understand this phenomenon, we may divide the equation 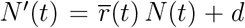 by *d*, leading to 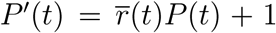 with *P* (*t*) = *N* (*t*)*/d*. Then, we observe that the main system (1) & (9) can be written in terms of *P* (*t*), independently of *N* and *d*. This means that the ratio *N* (*t*)*/d* is not influenced by *d*. This yields the independence of the evolutionary dynamics of *d*, because the effect of migration on mean fitness in (9) only depends on *d/N* (*t*).

A simple mathematical argument (Appendix F) shows that this property will apply beyond the present model. The result arises for any model where (i) the evolutionary and demographic dynamics in the sink are density-independent (apart from the impact of migration) and (ii) the sink is initially empty (or at least *d* ≫ *N* (0)). This means that it should apply for a broad class of models of asexual evolution in black-hole sinks. Note however, that sex and recombination, for example, necessarily create density-dependent evolution as recombination with migrants affects the genotype frequencies beyond the pure demographic impact of migration.

An intuition for the independence of 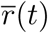 on *d* might be framed as follows: if *d* is increased (resp. decreased), the sink fills in more (resp. less) rapidly, from *N* (0) = 0, proportionally to the increase (resp. decrease) in *d*, at all times. Therefore things cancel out in the migration contribution on frequencies (*d/N* (*t*) is unaffected), and this contribution is the only one where *d* enters the dynamics. Overall increasing or decreasing *d* thus has no effect on genotype frequency dynamics, although it does affect population sizes. This balanced effect likely exists qualitatively in even more general conditions, but the exact cancelling out only happens with exponential (density-independent) growth/decay, density independent mutation and selection, and an initially empty sink.

#### Large time behavior

As seen in Fig. 2, 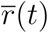 converges towards an asymptotic value 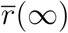 at large times. The expression (10) shows that this value depends on *r*_max_, *µ* and *n*. Interestingly, it becomes dependent on the harshness of stress *m*_*D*_, only in the case of establishment failure. More precisely, we get:

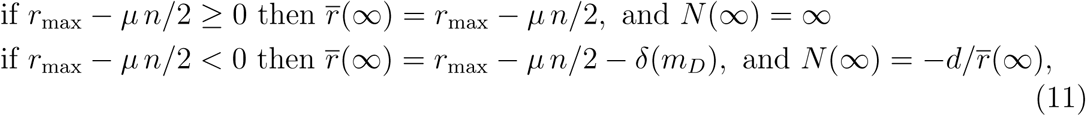

for some function *δ*(*m*_*D*_) such that *m*_*D*_ *> δ*(*m*_*D*_) *> m*_*D*_*/*8 for *µ* large enough (the inequality *δ*(*m*_*D*_) *> m*_*D*_*/*8 is true whatever the phenotype dimension *n*). When *n* is large enough, sharper lower bounds can be obtained, e.g. *δ*(*m*_*D*_) *>* 3 *m*_*D*_*/*8 for *n ≥* 6), see Appendix G.

These asymptotic results can be interpreted as follows. Below some threshold 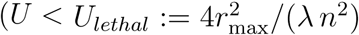, or equivalently *µ < µ*_*lethal*_ := 2*r*_max_*/n*), establishment is always successful and the sink population ultimately explodes (as we ignore density-dependence in the sink). As *d/N* (*∞*) = 0, the demographic and evolutionary effects of migrants thus become negligible (being diluted in an effectively infinite population). The sink population thus reaches mutation-selection balance, with a mutation load *µ n/*2, as if it was isolated. It ultimately grows exponentially at rate *r*_max_ − *µ n/*2 as illustrated in Fig. 2.

On the contrary, large mutation rates (*U ≥ U*_*lethal*_ or equivalently *µ ≥ µ*_*lethal*_) lead to establishment failure, which is a form of lethal mutagenesis (see Bull et al. (2007) for viruses and Bull and Wilke (2008) for bacteria) illustrated in Fig. 4. In this regime, the mutation load *µ n/*2 is larger than the absolute maximal fitness *r*_max_ in the sink. Therefore, at mutation-selection balance and even in the absence of any migration, the population could never show positive growth: establishment is impossible because the fitness peak is too low, given the mutation rate and effect. We further identify a “jump” of amplitude *δ*(*m*_*D*_) in the equilibrium mean fitness, as *µ* increases beyond the lethal mutagenesis threshold (illustrated in Fig. 5). Then, the population ultimately reaches a stable size determined by an immigration - decay equilibrium: a migration load can build up at equilibrium (*δ*(*m*_*D*_)) together with the mutation load (*µ n/*2). This migration load is produced by the constant inflow of maladapted genotypes from the source and does depend on the harshness of stress *m*_*D*_. It is this migration load that creates the “phase transition” in equilibrium fitness as *µ* crosses beyond *µ*_*lethal*_, the lethal mutagenesis threshold (Fig. 5). Note, however, that contrary to what happens with sexuals, migrants entering an asexual population do not interbreed with locally adapted genotypes, which simplifies the effect of migration. Note also that, in this lethal mutagenesis regime, the sink population does establish to a stable size, that may be higher than that expected in the absence of mutation and adaptation. However, this is not an establishment in that the population would still get extinct if migration was to be stopped.

### 3.2 Establishment time *t*_0_

Of critical importance is the waiting time until the sink becomes a source, when this happens, namely the time *t*_0_ at which 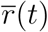 becomes positive. This section is devoted to the analysis of this time.

#### Derivation of an analytical expression

Using the expression (10), we can solve the equation 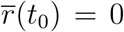. We recall that, due to our assumptions, *t*_0_ *>* 0, i.e.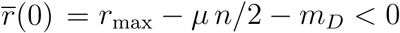.

**Figure 4:**
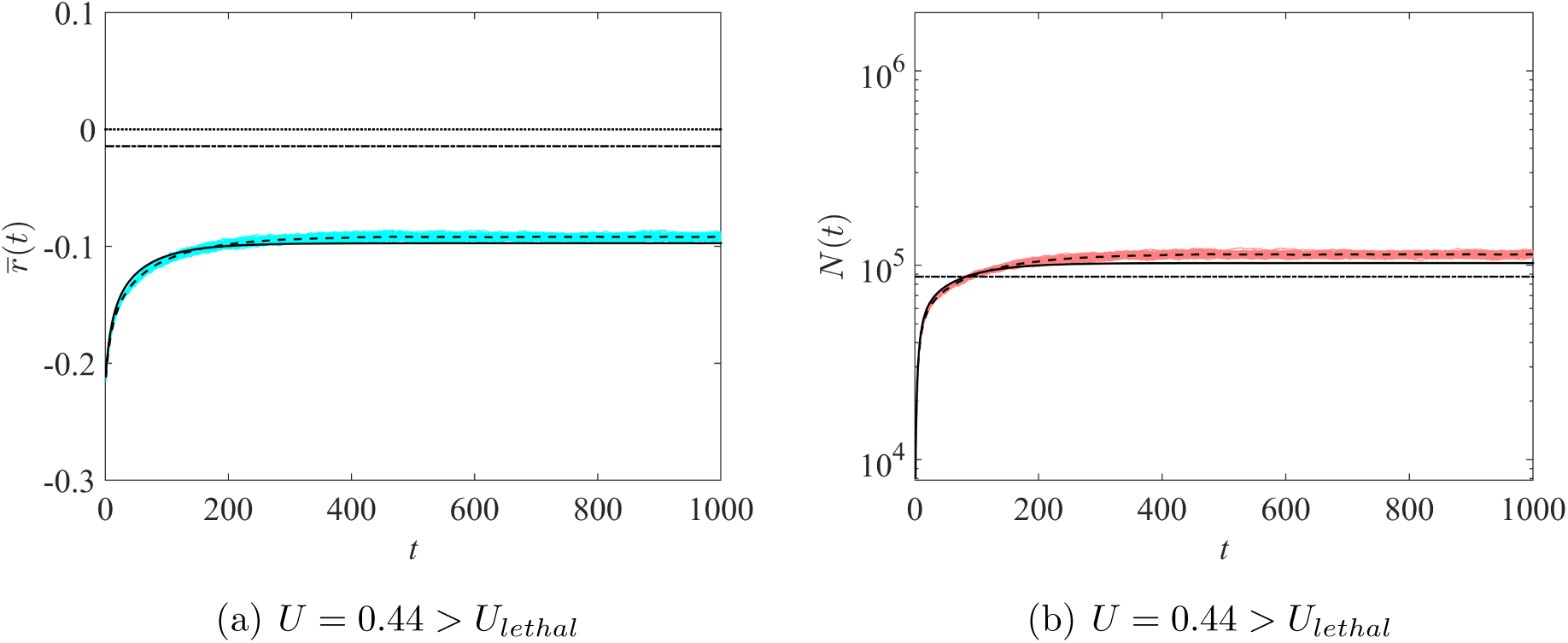
Trajectories of mean fitnesses and population sizes, lethal mutagenesis regime. Same legend as in Fig. 2. Other parameter values are *m*_*D*_ = 0.2, *r*_max_ = 0.1, *λ* = 1*/*300, *n* = 6 and *d* = 10^4^, leading to a theoretical threshold value for lethal mutagenesis 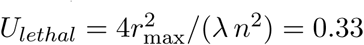. The panel (a) illustrates the bifurcation in the behavior of the equilibrium mean fitness as *r*_max_ − *µ n/*2 becomes negative.

**Figure 5:**
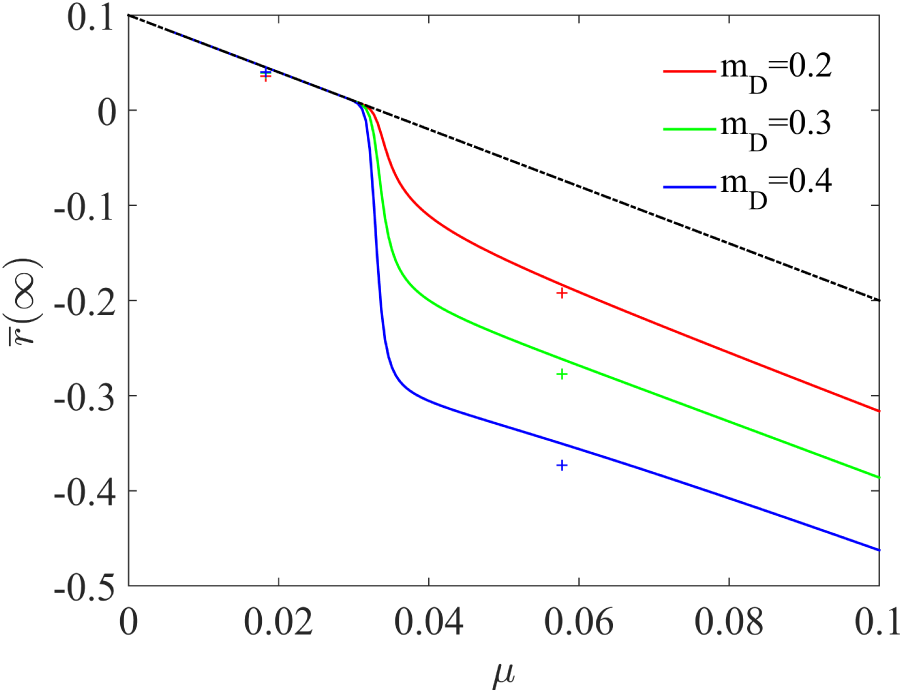
Mean fitness at large times, dependence with *µ* and *m*_*D*_. The solid lines are the values given by formula (11). The crosses correspond to the result of individual-based simulations. The dashed-dot line corresponds to *r*_max_ − *µ n/*2; the gap between the dashed-dot line and the solid lines represents the amplitude of the jump *δ*(*m*_*D*_). Parameter values: *r*_max_ = 0.1, *n* = 6.

The result in (11) shows that *t*_0_ = *∞* if *r*_max_ − *µ n/*2 *≤* 0 (establishment failure). In the case of successful establishment 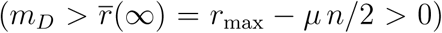, the waiting time to this establishment is:

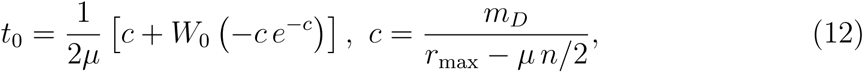

with *W*_0_ the principal branch of the Lambert-W function (see Appendix H).

First of all, eq. (12) shows that the waiting time is independent of the dispersal rate *d*. This was further supported by individual-based simulations (Fig. 6a) as *t*_0_ was found to drop rapidly to its predicted value as *d* increases (as the deterministic approximation becomes accurate), to then become independent of *d*. The waiting time shows a transition (around *c* = 1) from *t*_0_ *≈ c/*2*µ* for small *c* to *t*_0_ *≈ c/µ* for large *c*, so the establishment time always increases close to linearly with the harshness of stress *m*_*D*_. This was also the case in individual-based simulations (Fig. 6c), at least until stress becomes too strong, compared to mutation and migration. In that case, the sink population remains fairly small for a long time and our deterministic approximation no longer applies, at least in the early phases (1 and 2) of invasion (see Section 3.3). Eq. (12) also implies that the establishment time *t*_0_ decreases with *r*_max_ and increases with *n*. The dependence with respect to the mutational parameter *µ* is more subtle: as *µ* is increased, *t*_0_(*µ*) first decreases until *µ* reaches an ‘optimal value’ (minimizing invasion time), then *t*_0_(*µ*) increases until *µ* reaches the lethal mutagenesis threshold (*µ*_*lethal*_ = 2 *r*_max_*/n*). This behaviour always holds, as proven analytically in Appendix H. This non-monotonous variation of *t*_0_ with mutation rate (here with 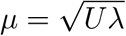) was also found in individual-based simulations (Fig. 6b).

Most of these effects are fairly intuitive: it takes more time to establish from a more maladapted source (*m*_*D*_), with a smaller mutational variance (*Uλ*), although their particularly simple quantitative effect on *t*_0_ was somewhat unexpected. The effect of *r*_max_, although quantitatively simple, has multiple aspects. Indeed, *r*_max_ affects various parameters of the establishment process, all else being equal: it decreases the initial rate of decay 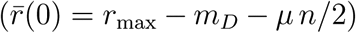 and increases the proportion of migrants that are resistant to the sink environment (fitness peak height) which both speed adaptation. It also increases the ultimate exponential growth rate of the population 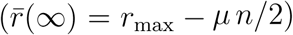. The latter effect is likely irrelevant to *t*_0_, however, as this growth phase occurs after the establishment time.

#### Effect of an intermediate sink

The simulations identify a sharp transition, in the harshness of stress, beyond which establishment does not occur (or occurs at very large times), see Appendix I. We see in Fig. 6 that as *m*_*D*_ gets close to this threshold, the dependence between *t*_0_ and *m*_*D*_ shifts from linear to superlinear (convex). Based on previous results on evolutionary rescue in the FGM (Anciaux et al., 2018), we conjecture that this pattern is inherent to the phenotype fitness landscape model. In the FGM, increased stress (higher *m*_*D*_) is caused by a larger shift in optimum from source to sink. This has two effects, (i) a demographic effect (faster decay of new migrants, on average) and (ii) an evolutionary effect. This latter effect is simply due to the geometry of the landscape. Indeed, when the shift in optimum from source to sink is larger, there are fewer genotypes, in the migrant pool, that can grow in the sink and they tend to grow more slowly. This effect is highly non-linear with stress, showing a sharp transition in the proportion of resistant genotypes beyond some threshold stress (for more details see Anciaux et al., 2018).

**Figure 6:**
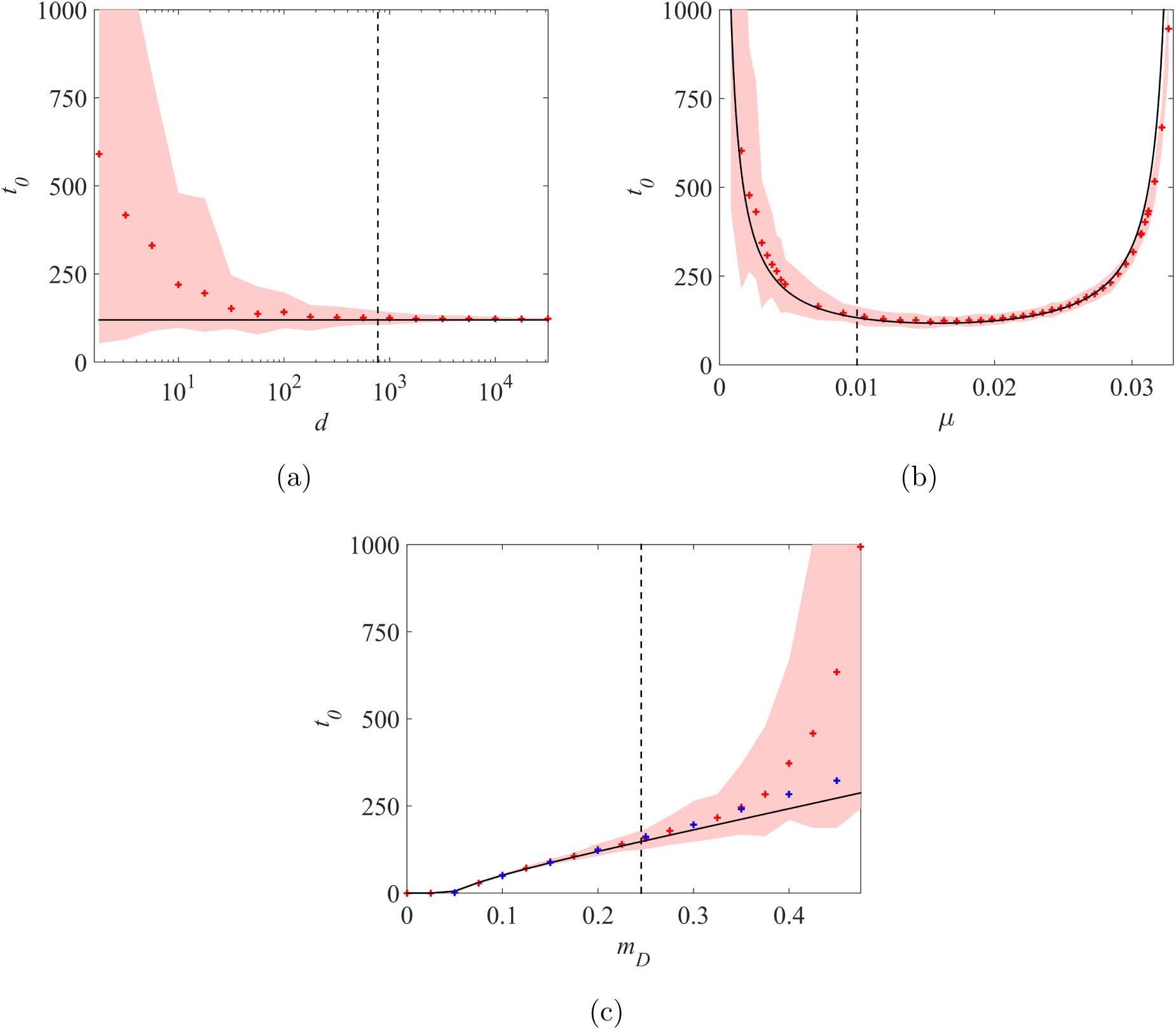
Establishment time *t*_0_, dependence with the immigration rate *d*, the mutational parameter *µ* and the harshness of stress *m*_*D*_. Theoretical value of *t*_0_ (black curve) vs value obtained with individual-based simulations (red crosses) and 95% confidence intervals, with fixed *m*_*D*_ = 0.2, *U* = 0.1 (panel a), *m*_*D*_ = 0.2, *d* = 10^3^ (panel b) and fixed *d* = 10^3^, *U* = 0.1 (panel c). The vertical dotted lines correspond to the values of *d, µ* and *m*_*D*_ such that 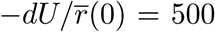 (panels b and c) and *U* = *U*_*c*_ (panel b). The blue crosses in panel (c) correspond to the establishment time 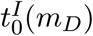, obtained by individual-based simulations, in the presence of an intermediate habitat with phenotype optimum **x**^*I*^ such that ‖**x***\* −* **x**^*I*^‖^2^*/*2 = ‖**x**^*I*^‖^2^*/*2 = *m*_*D*_*/*2. In all cases, the parameter values are *r*_max_ = 0.1, *λ* = 1*/*300, *n* = 6.

We argue that this type of dependence has important implications for the potential effect of an intermediate milder sink, with phenotype optimum **x**^*I*^ in between **x***\** (optimum in the source) and 0 (optimum in the sink), connected by a stepping-stone model of migration. A natural question is then whether the presence of this intermediate sink affects the waiting time to establish in the harsher sink. In that respect, assume that the overall harshness of stress (fitness distance between optima) is the same with and without the intermediate habitat *I*: schematically, *m*_*D*_ = *m*_*D*_(**x***\** → 0) = *m*_*D*_(**x***\** → **x**^*I*^)+ *m*_*D*_(**x**^*I*^ → 0). When *m*_*D*_ is low, *t*_0_ is roughly linear with *m*_*D*_ so that it may take a similar time to establish in two step and in one (the sum of intermediate establishment times would be the same as that to establish in a single jump). However, for harsher stress levels where *t*_0_ is superlinear with *m*_*D*_, the intermediate habitat could provide a springboard to invade the final sink, if both intermediate jumps are much faster than the leap from source to final sink.

To check this theory, we considered a new individual-based model with an intermediate habitat with phenotype optimum **x**^*I*^ such that ‖**x***\* −* **x**^*I*^‖^2^*/*2 = ‖**x**^*I*^‖^2^*/*2 = *m*_*D*_*/*2. The dynamics between the source and the sink are the same as those described in Section 2.5. In addition, we assume that (1) the source also sends migrants to the intermediate habitat at a rate *d*; (2) reproduction, selection and drift occur in the intermediate habitat following the same rules as in the sink, until the population *N*_*I*_(*t*) in the intermediate habitat reaches the carrying capacity *K* = *N \** (same population size as in the source); (3) the intermediate habitat sends migrants to the ultimate sink, at rate *d N*_*I*_(*t*)*/N \**. Then, we computed the time 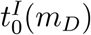 needed to establish in the final sink, in the presence of the intermediate habitat (value averaged over 100 replicate simulations).

The results presented in Fig. 6c (blue crosses) confirm that for small *m*_*D*_, the presence of an intermediate habitat has almost no effect 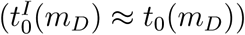. However, when *m*_*D*_ becomes larger and *t*_0_(*m*_*D*_) becomes superlinear, the establishment time in the sink is dramatically reduced by the presence of the intermediate sink 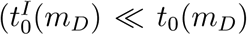; e.g., for 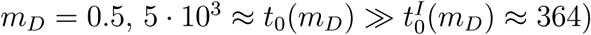.

#### Effect of mutation in the sink on the establishment time

We have seen in Fig 6b that mutation has a non-monotonous impact on establishment time. However, a higher mutation rate affects both the source equilibrium state and the sink dynamics. A natural question to ask is thus whether local mutation *in the sink* helps or hinders invasion. Indeed, mutation in the FGM (and other models with both deleterious and beneficial mutations) can have antagonistic effects: it generates fitness variance to fuel adaptation but lowers the mean fitness by creating a mutation load. This is of course also true for mutation in the source, but the interaction with migration in the sink makes the outcome less straightforward to grasp.

To tell apart the influences of local mutation on invasion speed, we analyzed (Appendix J) a scenario where mutation is absent in the sink, but still active in the source, so that the latter is unchanged. An expression equivalent to eq. (10) is obtained in this case for the mean fitness trajectory. We compared the corresponding time to establishment, noted 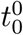, with the establishment time *t*_0_ to check whether local mutation (in the sink) speeds or slows invasion.

The results in Fig. 7 show that local mutation can either slow down or accelerate invasion, depending on the mutational variance (*µ*) and stress level (*m*_*D*_). For a given level of stress (*m*_*D*_), local mutation tends to speed invasion as long as mutational variance (*µ*) is limited (left part of the graph) but hinders it when it becomes larger (right part of the graph). The transition from helping to hindering invasion happens at larger *µ* values when the stress is harsher (higher *m*_*D*_). It thus appears that the beneficial effect of local mutation in producing variance dominates when mutation is limited while its negative effect in load buildup takes over as *µ* is increased. The transition occurs at higher *µ* under harsher stress because the former effect is more critical then, while the latter is roughly independent of stress. This pattern illustrates quite strikingly the complex implications, for adaptation dynamics, of the ambivalent nature of mutation in the FGM.

### 3.3 Range of validity of the model

**Figure 7:**
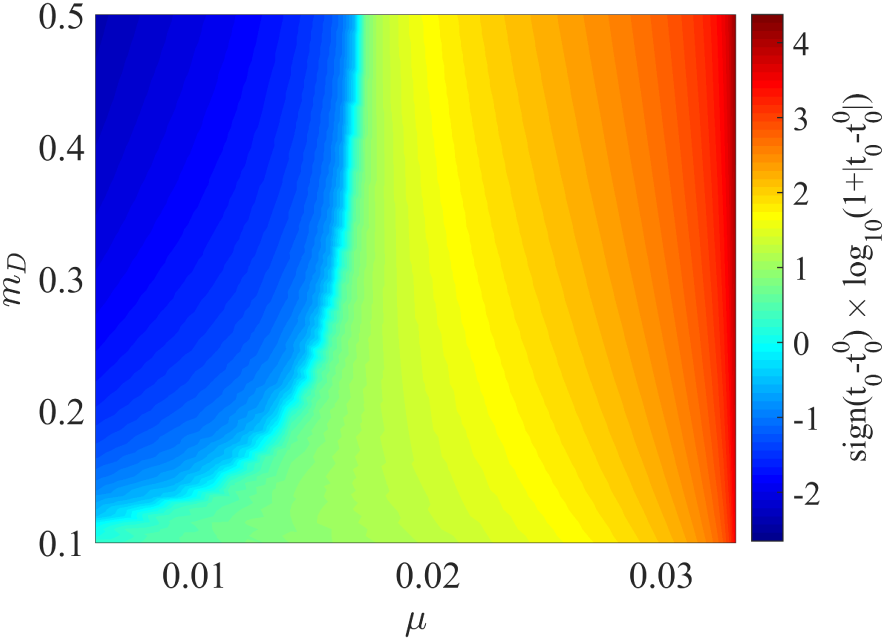
Comparison between the establishment times *t*_0_ (with mutation in the sink) and 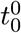 (without mutation in the sink). The heat map corresponds to sign 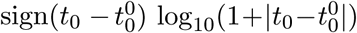: negative values indicate that 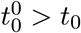 (faster establishment with mutation in the sink) and positive values indicate that 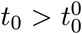 (faster establishment without mutation in the sink).

We explored the range of validity of the analytical model by comparing theory and simulations over a wide range of parameter values. The raw results are given in Appendix I. Overall, the model is more accurate as *U* and *d* increase and *m*_*D*_ (equivalently, *r*_*D*_ = *m*_*D*_ − *r*_max_), *n* and *λ* decrease. More precisely, theoretical and numerical analysis yield two (*a priori* conservative) conditions that should lead to the model being accurate: (i) *U ≥ U*_*c*_ = *n*^2^*λ/*4, for the WSSM to apply; (ii) *d U/r*_*D*_ ≫ 1, for the large *d* approximation to apply.

Below we detail each criterion, their robustness and possible empirical insight on their realism.

#### Criterion (i)

it is formally derived in Appendix E of (Martin and Roques, 2016) and guarantees that the mutation term associated with the FGM linearizes to produce an analytically tractable PDE. While the model is indeed accurate whenever *U > U*_*c*_, it remains reasonably so even at fairly lower mutation rates. Even for mutation rates *U* = *U*_*c*_*/*30 (but keeping a large *d*), 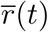 and *N* (*t*) from eq. (10) still accurately capture the average trajectories (Fig. 8), although the length of Phase 2 in the numerical simulations becomes more variable, around this average, as *U* is decreased. Consistently, Fig. 6b shows that the invasion time in eq. (12) accurately captures the average of simulations far below *U* = *U*_*c*_, with larger variability around this mean as *U* decreases.

As an example, empirical estimates in *E. coli*, based on a recent mutation accumulation experiment (Trindade et al., 2010) suggest *U ∈* [0.004, 0.006] and 𝔼[*s*] = *nλ/*2 *∈* [0.02, 0.04] (mean effect of mutations on fitness), which yields *U/U*_*c*_ *∈* [0.2, 0.6] for *n* = 1 and *U/U*_*c*_ *∈* [0.033, 0.1] for *n* = 6. This suggests that *E. coli* may lie somewhere below the critical mutation rate, at a similar order. Note however that estimates of these quantities are fairly scarce (even in this well studied biological model) and seem to vary substantially across experiments (medium, strain, growth conditions). We suspect that viruses (especially RNA viruses) may lie well within *U ≥ U*_*c*_, while bacteria may vary widely around *U* = *U*_*c*_. Obviously any proper statement on this issue would require a full review of empirical estimates (appropriately scaled in consistent time units), wherever available.

#### Criterion (ii)

this criterion, which is confirmed by the simulations in Appendix I and Fig. 6 panels (a) and (c), stems from the following argument: the early population size in the sink is of order 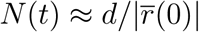 (no evolution), with 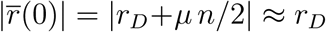 (when *µ ≪ r*_*D*_). Thus whenever *d* ≫ *r*_*D*_*/U,* the mutant input *N* (*t*)*U* in the sink population quickly reaches a large value *N* (*t*) *U ≈ d U/r*_*D*_ ≫ 1 and only increases later on. Adaptive evolution can then take place within the sink, in a way that is accurately captured by a deterministic approximation (see the dotted lines in Fig. 6). Conversely, when *d* is smaller and/or *r*_*D*_ is larger, the early population size in the sink is small, so that the deterministic approximation does not apply anymore. In this case, we see that the time *t*_0_ is much more variable, and increases on average with smaller *d* and larger *r*_*D*_ (or equivalently *m*_*D*_), see Fig. 6.

Empirically evaluating the criterion (ii) requires estimates of *d, U, r*_*D*_ on the same timescale (hours, days, generations) in a well defined sink. Such estimates should be possible from dedicated experiments controlling the immigration rate, in strains with known mutational parameters, and environmental stresses with well characterized demographic effect. They would greatly help our understanding of source-sink dynamics. However, to the best of our knowledge, they are not available to date.

## 4 Discussion

We derived an analytically tractable PDE-ODE framework describing evolutionary and demographic dynamics of asexuals in a source-sink system. Comparison with individual-based stochastic simulations shows that the approach is accurate in the WSSM regime (large mutation rates compared to mutation effects) and with a large migration rate, and seems robust to mild deviations from this regime. This approach reveals the typical shape of the trajectories of mean fitness and population sizes in a sink: (1) in the case of establishment failure, after a brief increase, the mean fitness remains stable at some negative level which depends on the harshness of stress; (2) in the case of successful establishment, this “plateau” is followed by a sudden increase in mean fitness up to the point where it becomes positive and the sink becomes a source. Note that here, we ignored density dependent effects in the sink, so that mean fitness ultimately converges towards an equilibrium that is independent of any migration effect, the latter being diluted into an exploding population.

The three first phases predicted by the model, for the case of successful establishment, are qualitatively observed in (Dennehy et al., 2010), an experimental study of invasion of a black-hole sink (an asexual bacteriophage shifting to a new bacterial host). The “host shift” scenario in their Fig. 3 corresponds roughly to our scenario with a population evolved on the native host sending migrants to a new host. The conditions may differ however as the population may not be initially at equilibrium in the native host at the onset of migration. Yet, the dynamics are qualitatively similar to those in our Fig. 2, although the time resolution in the data is too limited to claim or test any quantitative agreement. An extension of the present work could be to allow for non-equilibrium source populations, which can readily be handled by the PDE (9) (reformulating *ϕ*(*z*) = *ϕ*(*z, t*)). However, our analytical result on *t*_0_ does rely on an equilibrium source population. Note also that the four phases identified here are observed, in simulations, even in the low *d* or low *U* regimes where our analytical derivations can break down quantitatively. Therefore, while the model may provide qualitatively robust insight, quantitative analyses are necessary to really test its predictions. This would ideally include associated measures of decay rates *r*_*D*_, mutation rate *U* and ideally maximal possible growth rate *r*_max_, with a known immigration rate *d*.

Quite unexpectedly, the evolutionary dynamics (especially the waiting time *t*_0_ to establishment) do not depend on the immigration rate. This emerges mathematically from the fact that the evolutionary dynamics only depend on the population size through the ratio *N* (*t*)*/d* between the current population size and the immigration rate, this ratio itself remaining independent of *d*. This is confirmed by stochastic individual-based simulations (Fig. 6a): establishment time roughly decreases as 1*/d* when *d* is small but indeed stabilizes as *d* becomes larger. More precisely, the result on the independence of *t*_0_ with respect to *d* should always hold with an initially empty sink and when *d U/r*_*D*_ ≫ 1 (see Section 3.3). In this case, the mutant input in the sink population is always large enough to enable our deterministic framework to accurately capture the evolution in the sink. This result *a priori* extends to any model where evolution and demography are density-independent. However density dependent effects on demography or evolution (including sexual reproduction) might alter this outcome. Yet, we argue that purely demographic effects due to a finite carrying capacity in the sink environment should have limited impact on the conclusions of our model, up until establishment time (as long as *K* is large enough).

In a black-hole sink experiment Perron et al. (2007) studied the evolution of resistance to two lethal doses of antibiotics and their combinations in the bacterium *Pseudomonas aeruginosa* (also asexual). Their experiment differs from our scenario in that the sink populations were initially filled with many “naive” individuals (*N*_0_ ≫ 1, amounting to an initial large single immigration event). The authors did notice that immigration rate *d* affected population densities, but this is not directly a test of our model: our deterministic model also predicts that *N* (*t*) should depend on *d*, only the mean fitness and time to establishment do not.

The independence between *t*_0_ and *d* is counter-intuitive if we consider sink invasion as a repeated evolutionary rescue ‘experiment’. Indeed, the immigration process in the sink could also be seen as a Poisson process of incoming new lineages (from the source), each having a given probability *p*_*R*_ to yield a rescue in the future (in the absence of new immigration), hence to ultimately turn the sink into a source. This probability *p*_*R*_ can be computed from evolutionary rescue theory, with various flavours: see (Orr and Unckless, 2014) for a context-independent single allele rescue model or, in the case of the FGM, using results in (Anciaux et al., 2018). By basic properties of Poisson processes, the waiting time *t*_1_ to the first arrival, in the sink, of such a future rescue lineage should be exponential with mean 1*/*(*d p*_*R*_), thus decreasing as 1*/d*.

However, this waiting time is different from the one computed here. Our *t*_0_ denotes the time at which the mean fitness of the sink population becomes positive in the absence of immigration, hence the time at which the sink has truly become a source. The evolutionary rescue approach above computes the time *t*_1_ at which a lineage *ultimately* destined to produce a resistant genotype, enters the sink. This lineage may be very rare by *t* = *t*_1_, it may even not be resistant itself but only destined to produce a mutant offspring that will be. The time at which the sink will *de facto* be a positively growing source can thus be far later. A study and comparison of both waiting times is interesting and feasible, but beyond the scope of the present paper. This remark, however, has one key implication: migration may be stopped long before *t*_0_ and the sink may still ultimately become a source, with some probability (even if this will be ‘visible’ much later).

Some insight into the possible effects of management strategies, e.g. quarantine (*d*), lethal mutagenesis (*U*), prophylaxis (*m*_*D*_ and *r*_max_), can be developed from the results presented here.

Migration (propagule pressure) is considered an important determinant of the success of biological invasions in ecology (Von Holle and Simberloff, 2005; Lockwood et al., 2005). Consistently, it has been shown that the factors increasing potential contacts between human populations and an established animal pathogen or its host tend to increase the risk of emergence of infectious diseases (Morse, 2001). Under the ‘repeated rescue approach’ above, it is indeed expected that emergence risk should increase as 1/contact rate. However, the present work shows that the time at which this emergence will be *de facto* effective (visible) may be unaffected by this contact rate. This means that care must be taken in the criteria chosen to evaluate strategies, and between the minimization of emergence risk *vs.* emergence time.

The use of a chemical mutagen to avoid the adaptation of a microbial pathogen and the breakdown of drugs is grounded in lethal mutagenesis theory (Bull et al., 2007; Bull and Wilke, 2008). Our approach successfully captures the occurrence of this phenomenon: the establishment fails when the mutation rate *U* exceeds a certain threshold, which depends on *r*_max_, on the mutational variance *λ* and on the dimension of the phenotypic space. Additionally, once this threshold is reached, the equilibrium mean fitness ceases to depend linearly on the mutational parameter 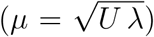, but rapidly decays (see Fig. 5). The existence of this negative “jump” in the equilibrium mean fitness, whose magnitude depends on the harshness of stress, leaves no room for evolutionary rescue. Conversely, our approach also reveals that below the lethal mutagenesis threshold, increasing the mutation rate decreases the establishment time as 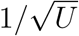. Hence, the use of a mutagen may be a double-edged sword since it can both hamper or increase the potential for adaptation in the sink.

As expected, the establishment time *t*_0_ increases with the harshness of stress *m*_*D*_; the population simply needs more time to adapt to more stressful environmental conditions. Increasing *m*_*D*_ or decreasing *r*_max_, whenever possible, are probably the safest ways to reduce the risks of biological invasions through adaptive processes or cross-species transmissions of pathogens (in both low and high *d* regimes). The precise dependence of *t*_0_ with respect to *m*_*D*_ brings us further valuable information. As long as our approach is valid (not too large stresses, leading to finite establishment times), a linear dependence emerges. It suggests that, in a more complex environment with a source and several neighbouring sinks connected by a stepping stone model of migrations, the exact pathway before establishment occurs in a given sink does not really matter. Only the sum of the stresses due to habitat shifts has an effect on the overall time needed to establish in the whole system. Conversely, for larger stress values our analytical approach is not valid, and the numerical simulations indicate a convex (surlinear) dependence of *t*_0_ with respect to *m*_*D*_. In such case, for a fixed value of the cumulated stress, the establishment time in the sink could be drastically reduced by the presence of intermediate sink habitats.

This result, which needs to be confirmed by more realistic modelling approaches and empirical testing, might have applications in understanding the role of so-called “preadaptation” in biological invasions. Recent adaptation to one or more facets of the environment within the native range has been proposed as a factor facilitating invasions to similar environments (e.g. Hufbauer et al., 2012, anthropogenically induced adaptation to invade). Our results suggest that preadaptation might only reduces the overall time to invasion (i.e., taking the preadapation period into account) only when invading highly stressful habitats.

The effect of a given environmental challenge, and thus their joint effects when combined (Rex Consortium, 2013), might be modelled in various ways in a fitness landscape framework (see also discussions in Harmand et al., 2017; Anciaux et al., 2018). The first natural option is to consider that multiple stresses tend to pull the optimum further away, and possibly lower the fitness peak *r*_max_. In the simplified isotropic model studied here, a larger shift in optimum amounts to increasing *m*_*D*_. However, a possibly more realistic anisotropic version, with some directions favored by mutation or selection, might lead to directional effects (where two optima at the same distance are not equally easy to reach) and be particularly relevant to multiple stress scenarios. Such a more complex model could be handled by focusing on a single dominant direction (discussed in Anciaux et al., 2018), or by following multiple fitness components (one per direction, Hamel et al. in prep).

Clearly, many developments are possible and could prove useful to understand how qualitative and quantitative aspects of environmental stresses may affect rescue and invasion. The present isotropic approach provides a simple, tractable null model for the latter, where all environmental effects are summarized by their measurable effects on *m*_*D*_, *U λ* and *r*_max_. We hope it will foster the empirical study of source-sinks with associated measurements of these key parameters.

## Acknowledgments

This preprint has been reviewed and recommended by Peer Community In Evolutionary Biology (https://doi.org/10.24072/pci.evolbiol.100072). This work was supported by the French Ministry of Higher Education, Research and Innovation (MESRI allocation doctorale to Y.A.), and the French Agence Nationale de la Recherche (ANR-13-ADAP-0016 “Silentadapt” to G.M., ANR-13-ADAP-0006 “MeCC” and ANR-14-CE25-0013 “NONLOCAL” to L.R. and ANR-18-CE45-0019 “RESISTE” to G.M. and L.R.). This work was fostered by stimulating discussions with Ophélie Ronce and François Hamel.

## A Fitness distribution of the migrants:derivation of formulae (5) and (6)

Consider an individual with phenotype **x**. Its fitness in the source is *m*_*source*_ = −‖**x** − **x**^⋆^‖^2^*/*2, where **x**^⋆^ is the optimal phenotype in the source, whereas its fitness in the sink is *m*_*migr*_ = −‖**x**‖^2^*/*2. We observe that

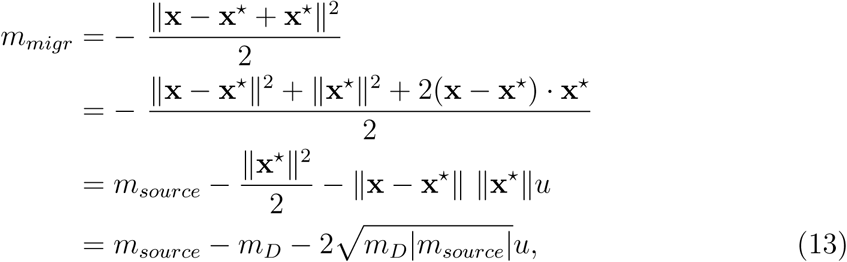

with *m*_*D*_ = ‖**x**^*\**^‖^2^*/*2 and a constant *u ∈* [−1, 1]. As the source is assumed to be at the mutation-selection equilibrium, the distribution of fitness in the source satisfies *m*_*source*_ *∼ −*Γ(*n/*2, *µ*) (Martin and Roques, 2016, equation (10)) and the corresponding moment generating function is 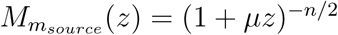. The results in (Martin and Lenormand, 2015) show that *u* is a random variable with moment generating function:

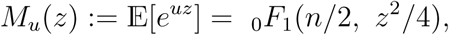

With _0_*F*_1_ the hypergeometric function, defined by 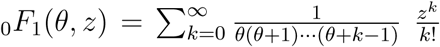.Let us first compute the moment generating function 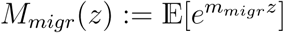. We have

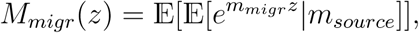

and using (13),

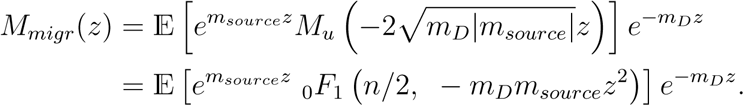

Thanks to the definition of the hypergeometric function _0_*F*_1_(*n/*2, *z*), we get:

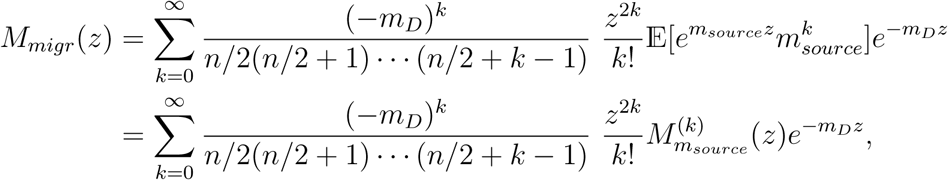

with 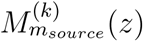 the k^th^ derivative of 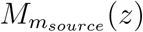 with respect to *z*. Thus,

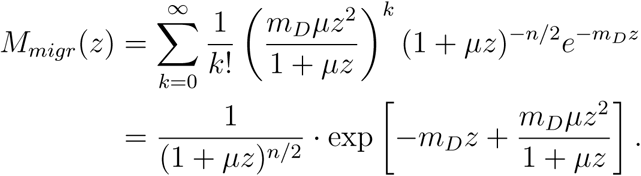

Setting *ϕ*(*z*) = ln (*M*_*migr*_(*z*)), we obtain formula (5).

Let us now show that the distribution of the migrants in the sink satisfies (6). Let *p*_*migr*_ be defined by (6). We just have to check that the moment generating function of *p*_*migr*_ is *M*_*migr*_:

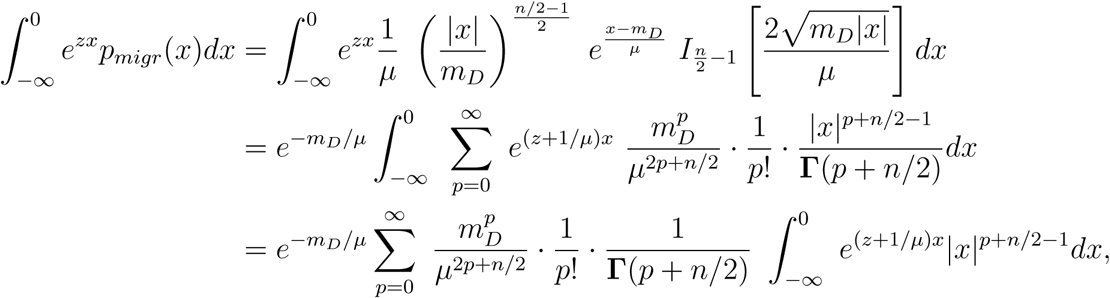

where *I*_*?*_ is the modified Bessel function of the first kind and **Γ** the gamma function.

Now, for all positive numbers *a* and *b*, we have:

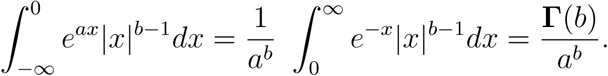

Therefore, we get, for *z > −*1*/µ*:

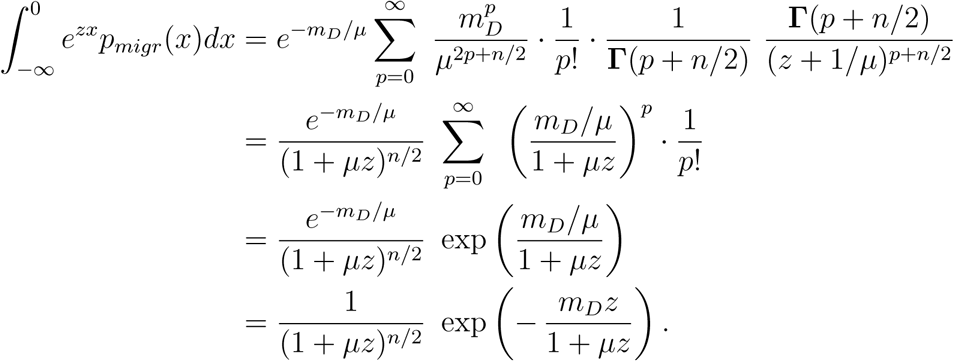

This is consistent with formula (5), which proves that the expression (6) is correct.

### B PDE satisfied by the CGF of the fitness distribution

In the WSSM regime, and in the absence of immigration, Martin and Roques (2016) (see Appendix E, equation (E5)) have shown that the CGF of the fitness distribution satisfies the following equation:

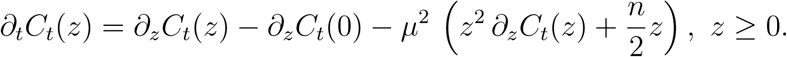

We derive here the additional term in (9), which describes the effect of immigration on the CGF.

In that respect, we consider a discrete population of size *N* (*t*) *∈* ℕ at time *t*, and the corresponding fitnesses (*m*_1_(*t*), *…, m*_*N*(*t*)_(*t*)). We define the “empirical” moment generating function

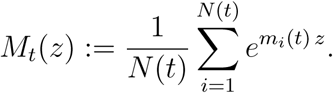

Assuming a Poisson number of immigration events, with rate *d* per unit time (see Section 2.5), for Δ*t* small enough, the probability that a single immigration events occurs during (*t, t*+Δ*t*) is approximately *d* Δ*t*. The probability that several immigration events occur during this time interval is close to 0. Therefore, the expected change in the moment generating function during Δ*t*, conditionally on the fitness *m*_*migr*_ of the unique migrant, is:

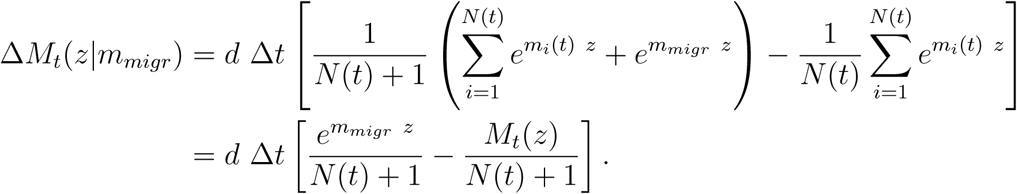

Taking expectation over the distribution of *m*_*migr*_ (see Appendix A for more details on the distribution of *m*_*migr*_), we get

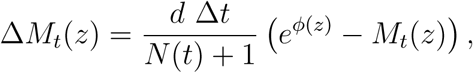

with 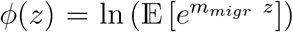. The corresponding change in the CGF *C*_*t*_(*z*) = ln *M*_*t*_(*z*) is Δ*C*_*t*_(*z*) *≈* Δ*M*_*t*_(*z*)*/M*_*t*_(*z*). Thus,

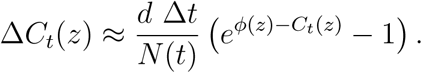

Finally, dividing the above expression by Δ*t* and passing to the limit Δ*t* → 0, we obtain the last term in (9), which describes the effect of immigration on the CGF:

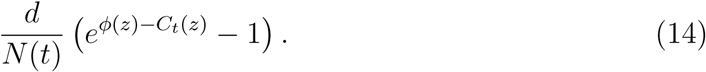

### C Solution of the system (1) & (9)

This section is devoted to the mathematical study of the system (1) & (9). We rewrite it in the following form:

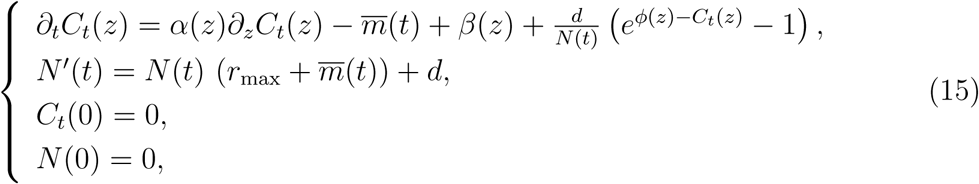

with *t >* 0 and *z ≥* 0, and where *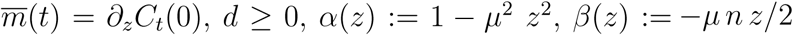.*

We can easily check that the sink is not empty at each time *t >* 0:

#### Lemma 1.

*Assume that* 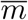 *is continuous over* [0, *∞*). *Then, at all time t >* 0, *we have N* (*t*) *>* 0.

*Proof.* For *ε >* 0 small enough, as *N′* (0) = *d >* 0, we have *N* (*t*) *>* 0 for all *t ∈* (0, *ε*]. Additionally, for all *t ≥ ε*, ?

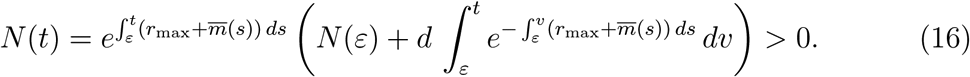

Let *N* (*t*), *C*_*t*_(*z*) be a solution of (15), such that 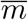 is continuous over [0, *∞*). Set *D*_*t*_(*z*) = *C*_*t*_(*y*(*z*)), with *y*(*z*) = tanh(*µz*)*/µ* which satisfies:

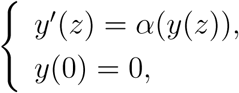

so that

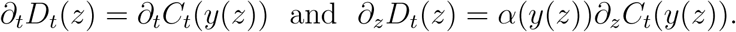

Thus, *D*_*t*_(*z*) satisfies the simpler equation

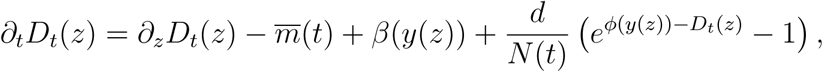

with 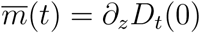.

Using the method of characteristics, we derive an analytic expression for *D*_*t*_(*z*). Fix *z ≥* 0 and denote for all *z ≥ t >* 0:

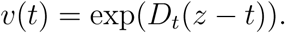

The function *v ∈ C*^1^((0, *z*]) satisfies for all *t ∈* (0, *z*):

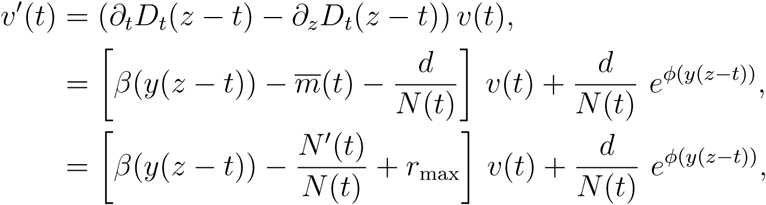

thanks to 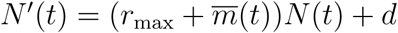. Let us fix times 0 *< ε < t*. By Lemma 1, we know that *N* (*s*) *>* 0, for all *s ∈* [*ε, t*] and so *v*(*t*) is given by:

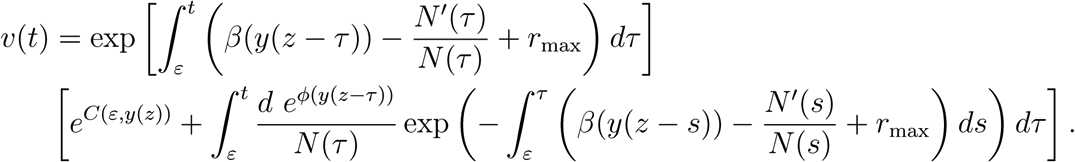

As 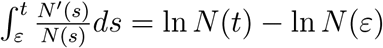, we can simplify the last expression to:

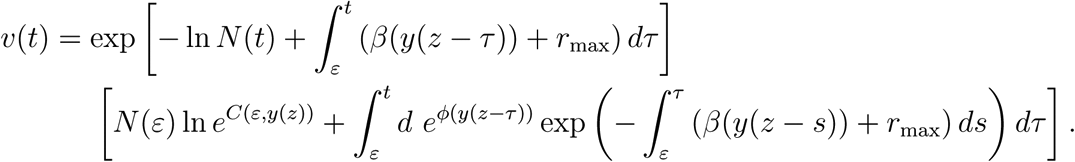

Taking the limit as *ε* tends to 0 and using the fact that the initial population in the sink is *N* (0) = 0, the above expression can be simplified to:

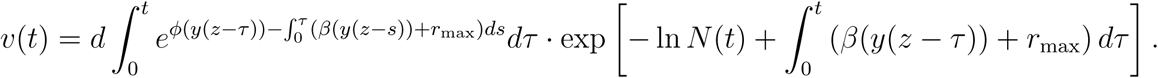

Hence, by reversing the characteristics, we get:

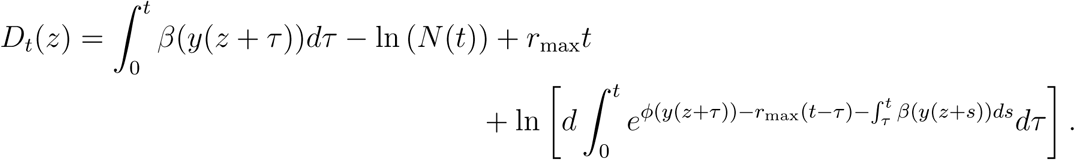

This leads to an explicit but complex formula for *C*_*t*_(*z*) thanks to the relation

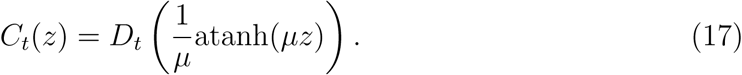

Additionally, we have:

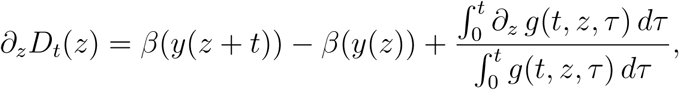

with 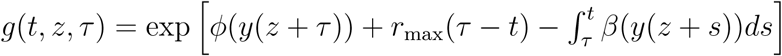. Using the fact that 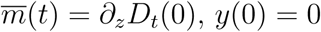 and *β*(0) = 0, we get:

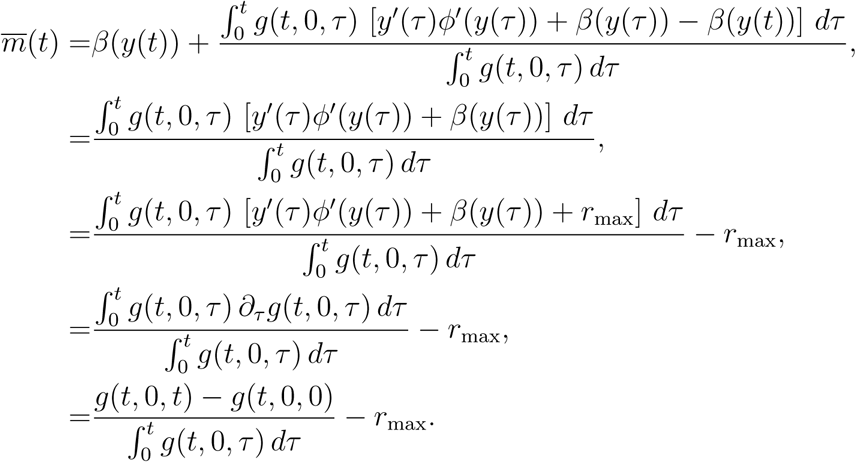

Using the expression 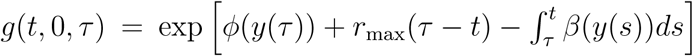, the formula (5) for *ϕ* and *y*(*z*) = tanh (*µz*)*/µ*, we finally get:

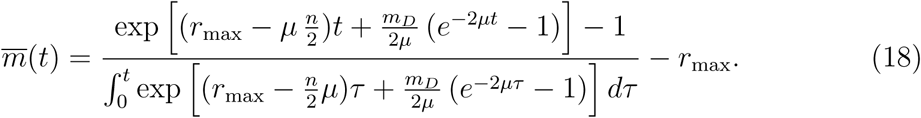

As we have an explicit formula for *m*(*t*), we can also solve the ODE 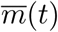 (formula (16), with *ε* = 0 and *N* (*ε*) = 0). Finally, we can check that *N* (*t*), *C*_*t*_(*z*) (defined by (17)) is a solution of (15) such that 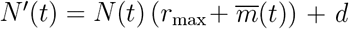 (given by (18)) is continuous over [0, *∞*). Using the expression (18) with 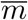, we obtain the formula (10) in the main text.

### D Trajectories of mean fitness: *U < U*_*c*_

**Figure 8:**
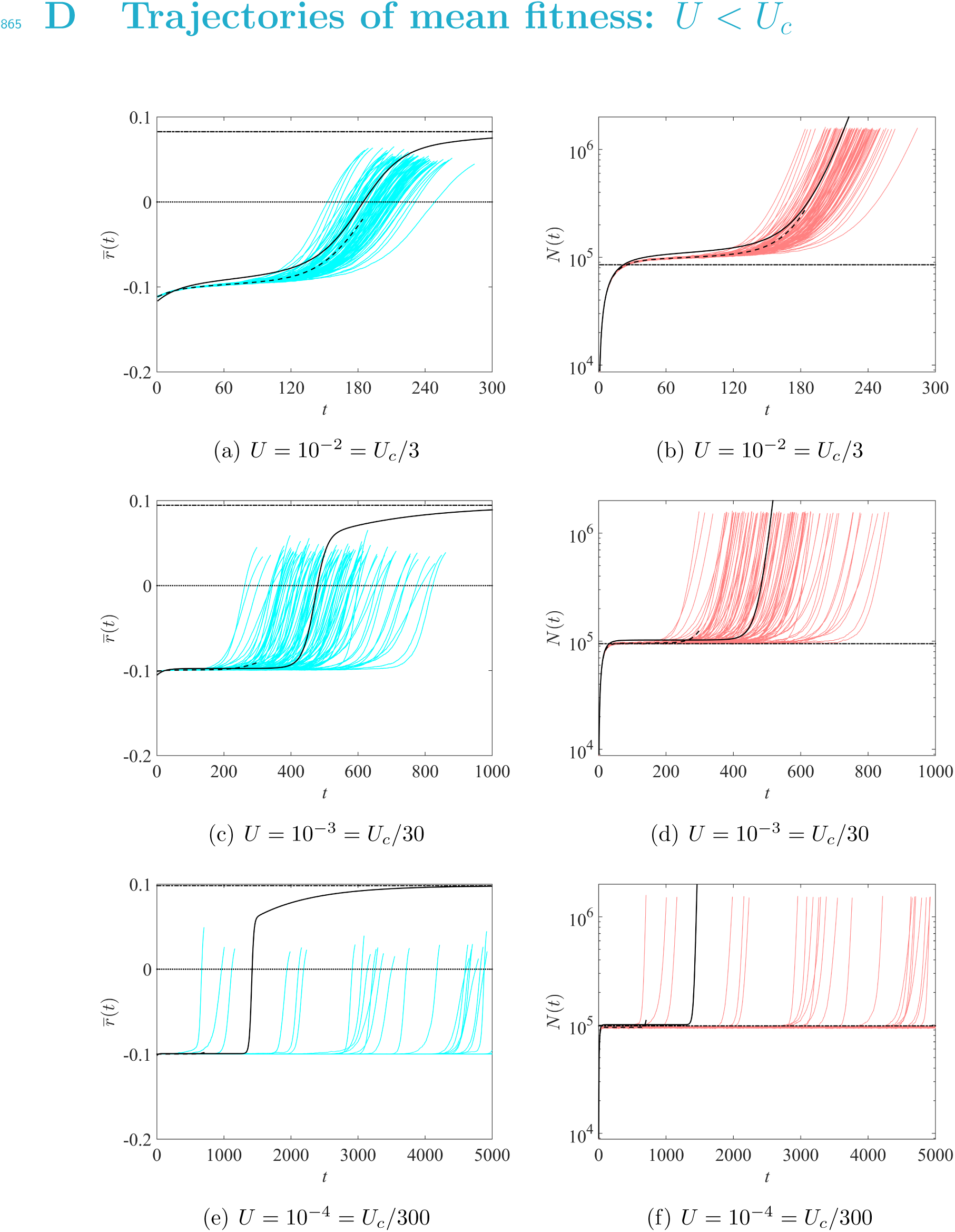
Trajectories of mean fitnesses and population sizes, low mutation rates. Same legend as in Fig. 2. Other parameter values are *m*_*D*_ = 0.2, *r*_max_ = 0.1, *λ* = 1*/*300, *n* = 6 and *d* = 10^4^, leading to *U*_*c*_ = 0.03.

### E Phenotype distribution in the sink: dynamics of 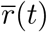 and *N* (*t*)

The dynamics of mean fitness and population size corresponding to Fig. 3 are plotted in Fig. 9, to illustrate the occurrence of the four phases in this particular simulation.

**Figure 9:**
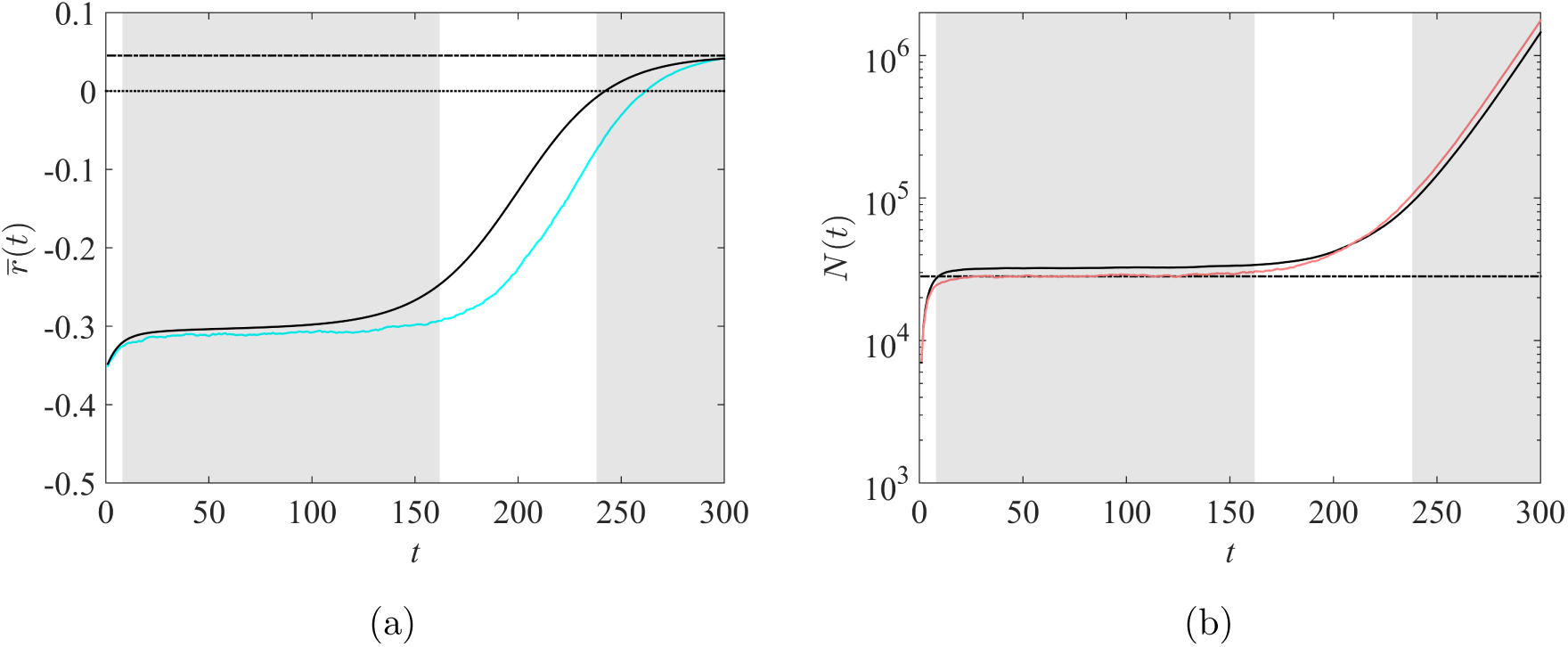
Trajectory of mean fitness and population size in the sink corresponding to the phenotype distribution in Fig. 3. Same legend as in Fig. 2.

### F Independence of the evolutionary dynamics with respect to the immigration rate

The value of 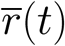 in formula (10) does not depend on *d*. Thus, only the population size dynamics are influenced by the immigration rate, but not the dynamics of adaptation. Actually, this phenomenon appears for a more general deterministic black-hole sink model, with a stable source and a constant immigration rate *d ≥* 0. In the sink, we have just to assume that the environment is initially empty (*N* (0) = 0), that both demography and evolution are density-independent (so that density dependence only arises in the migration effect). Apart from that, the proposed generalization may accommodate arbitrary forms of mutation and selection effects (possibly with changes in stress over time). The model then takes the following general form:

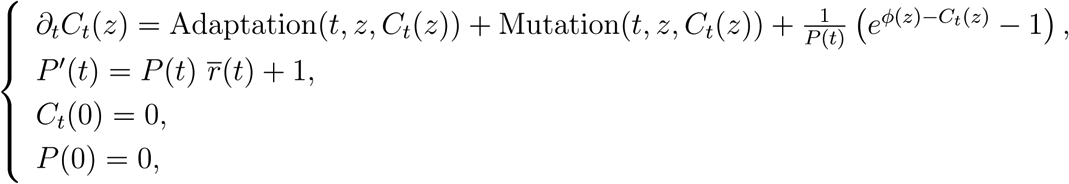

with 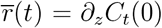 the coefficient of the exponential growth. Setting *P* (*t*) = *N* (*t*)*/d*, we observe that the above system can be written in the form:

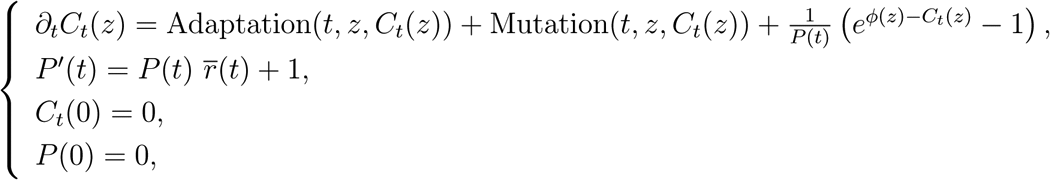

with 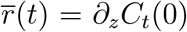. As this system does not depend on *d*, this implies that the dynamics of *P* (*t*), of mean fitness 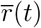, and even of the full fitness distribution (*C*_*t*_(*z*)) are all independent of *d*.

### G Large time behavior of 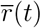

We recall that, according to formula (10),

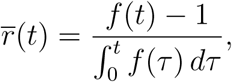

with 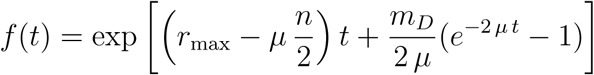.

*We first show that 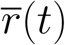 is an increasing function of t.* First, we can check that

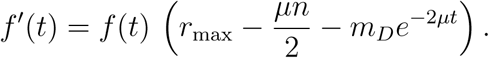

Second, we have

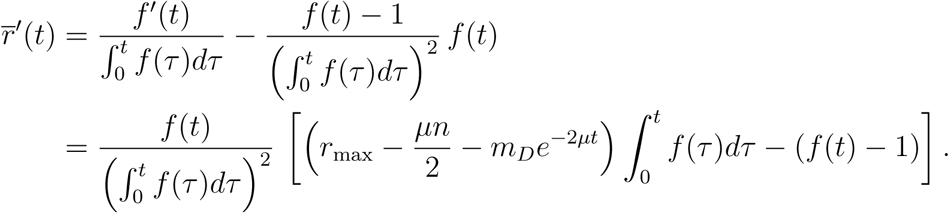

Let 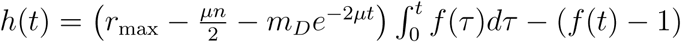. Thus we see that

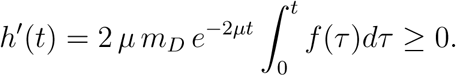

Therefore for all *t >* 0, *h*(*t*) *> h*(0) = 0, which shows that 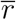 is increasing.

Since 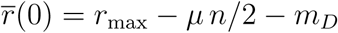, this implies that 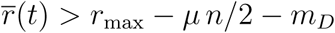 for all *t >* 0. In particular, 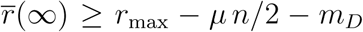 which implies that *δ*(*m*_*D*_) *< m*_*D*_ in (11).

*Next, we compute the limit of 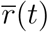 as t* → *∞.*

Case (i): we assume that *r*_max_ − *µn/*2 *>* 0. Then, *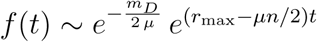* and

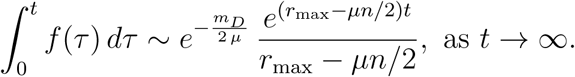

Thus,

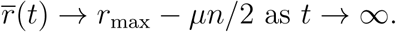

Case (ii): we assume that *r*_max_ − *µn/*2 = 0. Then 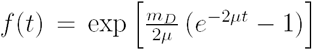 and 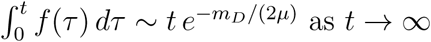 as *t* → *∞*. Thus,

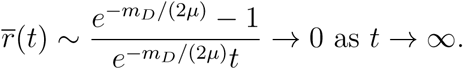

Case (iii): we assume that *r*_max_ − *µn/*2 *<* 0. Consider an arbitrary constant *α ∈* (0, 2). We can check that, for all 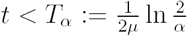, we have:

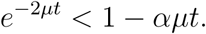

In the sequel, we denote *X* := *r*_max_ − *µn/*2. We get:

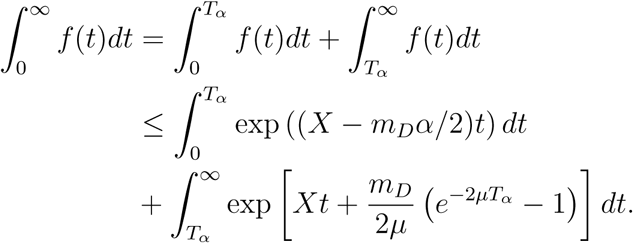

Using the assumption *X* = *r*_max_ − *µn/*2 *<* 0, we obtain:

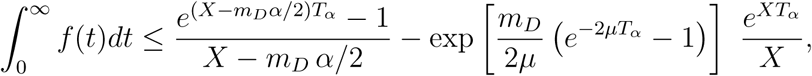

and using the definition of 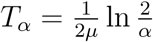, we obtain

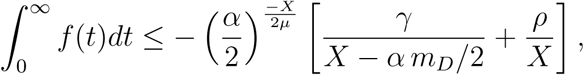

with 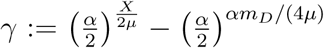 and 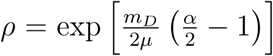. This leads to the following inequality:

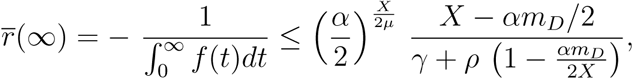

which can be rewritten:

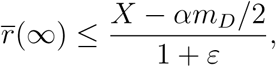

with

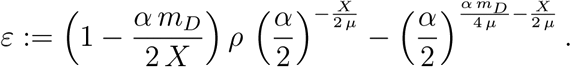

Next, to show that 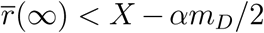, we only need to check that *ε <* 0. This is true for certain values of *α*. As 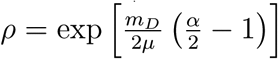, we observe that *ε* has the same sign as:

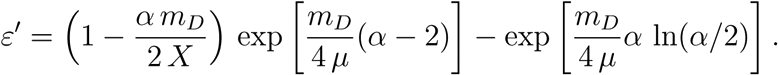

Since *X* = *r*_max_ − *µn/*2, we get:

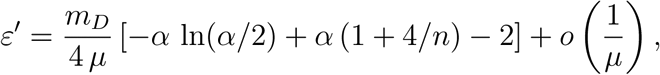

as *µ* → *∞*. Thus, *ε <* 0 for *µ* large enough, if and only if:

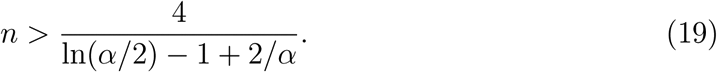

For *α* small enough, this inequality is true for any *n ≥* 1. However, higher values of *α* lead to sharper estimates of *δ*(*m*_*D*_) in (11). With *α* = 1*/*4 for instance, the inequality (19) is always satisfied (as *n ≥* 1). We obtain that 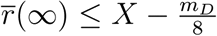 and 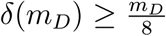 for *µ* large enough. If *α* is increased, e.g., *α* = 1*/*2, the inequality (19) is true for all *n ≥* 3, and consequently, 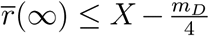 for *µ* large enough (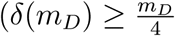, for *µ* large enough). In our numerical computations (*n* = 6), we can use *α* = 3*/*4, which leads to 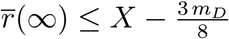 and 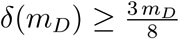 for large *µ*.

### H Establishment time *t*_0_: formula (12)

We recall that *t*_0_ is defined as the first zero of 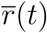. We note that, since 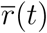 is increasing, it admits at most one zero.

First, if *r*_max_ − *µn/*2 *≤* 0, as 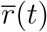 is increasing and 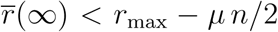 (see (11) and Appendix G), we have 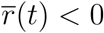 for all *t ≥* 0. This implies that *t*_0_ = *∞*.

Second we assume that *r*_max_ − *µn/*2 *>* 0. In this case, 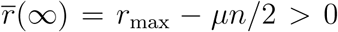 and the time *t*_0_ is finite (and positive). Therefore, we can solve the equation 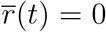, which is equivalent to:

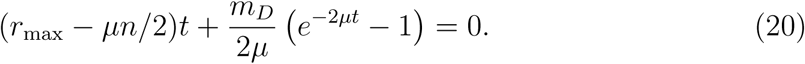

Let us set *c* := *m*_*D*_*/*(*r*_max_ − *µ n/*2). Since 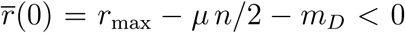, we observe that *c >* 1. The equation (20) is equivalent to:

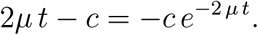

Multiplying this expression by *e*^2*µ t-c*^, we get:

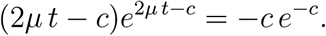

Setting *X* := 2*µ t - c*, we obtain:

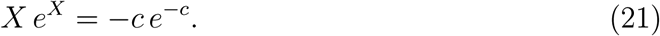

As *c >* 1, *-ce*^*-c*^ *∈* (*-e*^−1^, 0), thus the equation (21) admits two solutions, *X*_0_ = *W*_0_(*-c e*^*-c*^) and *X*_−1_ = *W*_−1_(*-c e*^*-c*^) *< X*_0_, with *W*_0_ and *W*_−1_ respectively the principal branch and the lower branch of the Lambert-W function. Thus, the equation (20) admits two solutions, (*c* + *X*_0_)*/*(2*µ*) and (*c* + *X*_−1_)*/*(2*µ*) = 0, but only the first one is positive. Finally, we obtain that

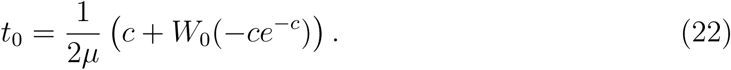

As *t*_0_ is an increasing function of *c*, we obtain that *t*_0_ decreases as *r*_max_ is increased, and *t*_0_ increases as *m*_*D*_ and *n* are increased. The dependence with respect to *µ* is more subtle. Differentiating the expression (22) with respect to *µ*, we observe that 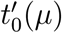 has the same sign as:

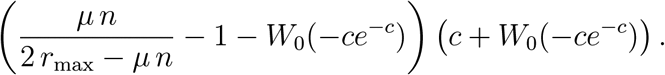

As the second factor in the above expression is always positive (since *c >* 1), we get that 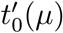 has the same sign as the function:

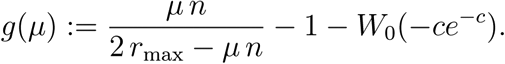

Differentiating *g* with respect to *µ*, we observe that *g′* (*µ*) has the same sign as *r*_max_ + (*µ n/*2 + *m*_*D*_) *W*_0_(*-ce*^*-c*^) = *r*_max_ − *µ n/*2 + *µ n* (1 − *W*_0_(*-ce*^*-c*^))*/*2 + *m*_*D*_ *W*_0_(*-ce*^*-c*^). Thus *g′* (*µ*) has the same sign as *m*_*D*_ (1*/c* + *W*_0_(*-ce*^*-c*^)) + *µ n*(1 − *W*_0_(*-ce*^*-c*^))*/*2 *>* 0, as 1*/c* + *W*_0_(*-ce*^*-c*^) *>* 0 (since *c >* 1) and 1 − *W*_0_(*-ce*^*-c*^) *>* 0. Finally, *g* is increasing, with:

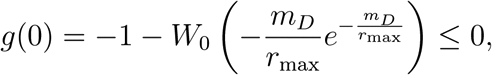

(and the sign is strict unless *m*_*D*_ = *r*_max_). Additionally, we have *g*(2*r*_max_*/n*) = +*∞* (corresponding to *µ*_*lethal*_). This means that, unless *m*_*D*_ = *r*_max_, *t*_0_(*µ*) first decreases until *µ* reaches an optimal value, and then increases as *µ* is increased.

### I Establishment time *t*_0_: dependence with the harshness of stress *m*_*D*_ and the immigration rate *d*

Using the stochastic individual-based model of Section 2.5, we analysed the dependence of the establishment time *t*_0_ with respect to *m*_*D*_ and *d* for a wide range of parameter values. Namely, taking *U* = 0.1, *r*_max_ = 0.1, *λ* = 1*/*300 and *n* = 6 as in Fig. 6, *m*_*D*_ was varied between 0.1 and 0.5. The results are presented in Fig. 10a. It shows that, for each value of *m*_*D*_, there is a threshold value of the immigration rate above which the establishment time *t*_0_ becomes almost independent of *d*. This threshold tends to increase as the harshness of stress *m*_*D*_ takes higher values. Additionally, we measured the relative error between the theoretical value of *t*_0_ given by formula (12) and the value given by individual-based simulations; see Fig. 10b. As soon as the parameters are far from the black region in Fig. 10, (a,b), the approximation is accurate (relative error *<* 0.1). This black region corresponds to values of *t*_0_ *>* 5000, for which individual-based simulations were stopped before establishment, and where we can expect that the final outcome is establishment failure. This means that there is only a narrow region where formula (12) is not accurate; it is located close to the region where establishment fails, and describes a rapid increase in *t*_0_ which is not captured by our analytical approach. Fig. 10 (c,d) depicts comparable simulations, with *U* = *U*_*c*_*/*3 = 0.01, i.e., outside of the WSSM regime. The conclusions are similar to the case *U* = 0.1, but with a larger region corresponding to establishment failure, and a lower accuracy (panel d).

**Figure 10:**
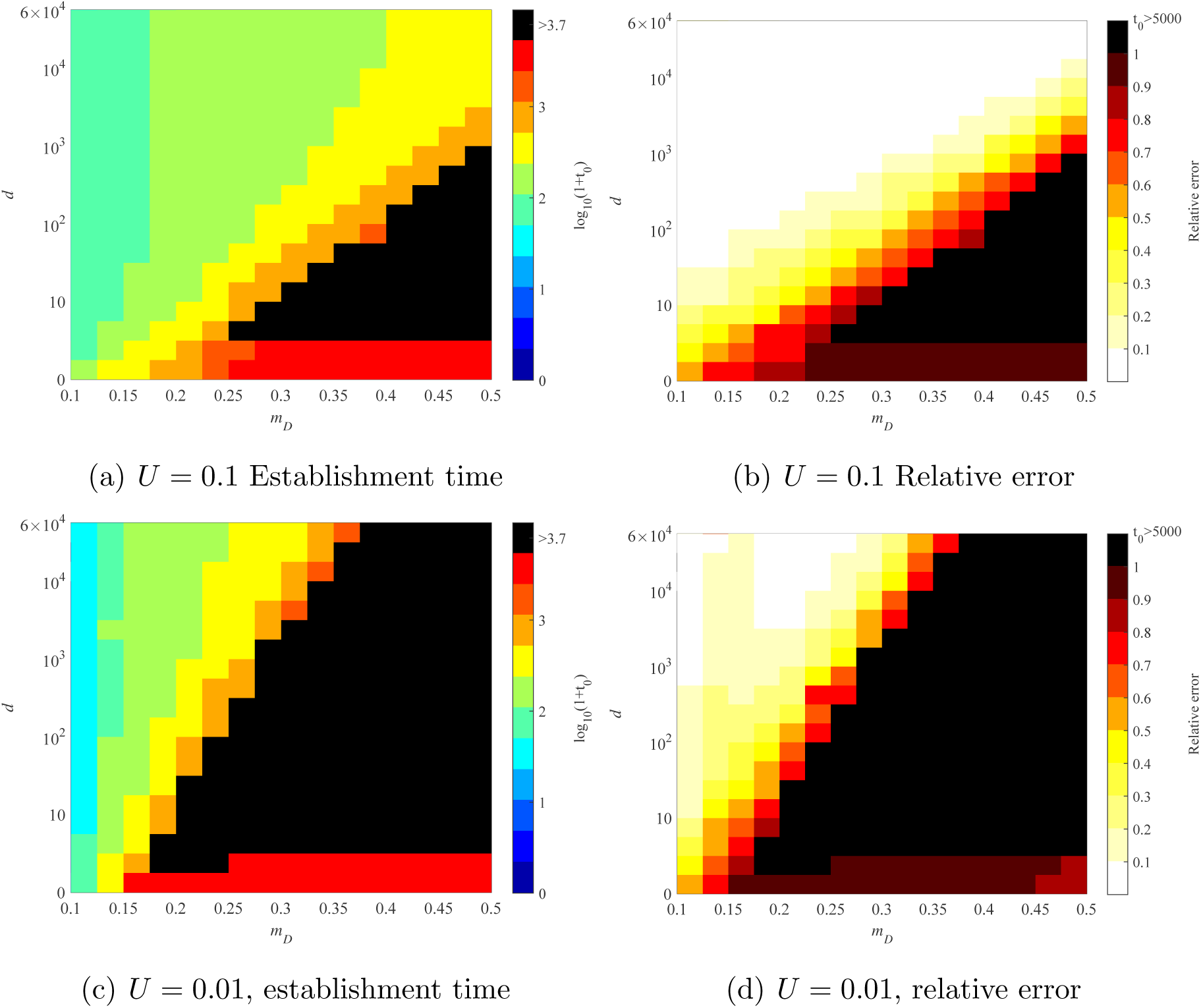
Establishment time *t*_0_, dependence with the harshness of stress *m*_*D*_ and the immigration rate *d*. (a,c): Average value of *t*_0_ over 100 individual-based simulations. The color legend corresponds to log10(1+*t*_0_). (b,d): relative error between the theoretical value of *t*_0_ given by formula (12) and the average value obtained by individual-based simulations. The black regions correspond to parameter values for which at least one simulation led to *t*_0_ *>* 5000; in that case, the average value of *t*_0_ was not computed numerically. In all cases, the parameter values are *r*_max_ = 0.1, *λ* = 1*/*300, *n* = 6.

### J Dynamics in the absence of mutation in the sink

To get a better understanding of the four phases described in Section 3.1, we considered the case where the mutation rate *U* = 0 in the sink (while it remains positive in the source).

First, using the same arguments as in Appendix C, we can derive a formula for 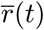 in that case. The formula can be expressed in the same form as (10), with:

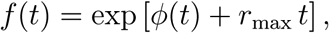

with *ϕ* given by (5).

An example of trajectory of fitness is given in Fig. 11, where we observe that the four phases are still present. The corresponding phenotype distribution is presented in Fig. 12. A video file of the phenotype distribution is also available as Supplementary File 3.

**Figure 11:**
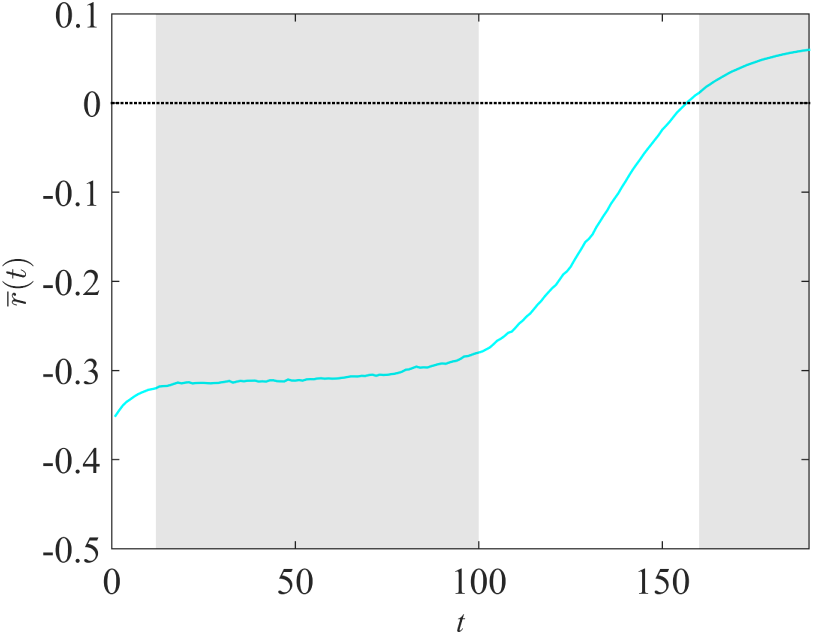
Dynamics of 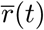 in the absence of mutation in the sink. The blue curve corresponds to the trajectory of 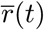 given by a single individual-based simulation, in the absence of mutation in the sink. The parameter values are *m*_*D*_ = 0.4, *U* = 0.1, *r*_max_ = 0.1, *λ* = 1*/*300, *n* = 6 and *d* = 10^4^.

**Figure 12:**
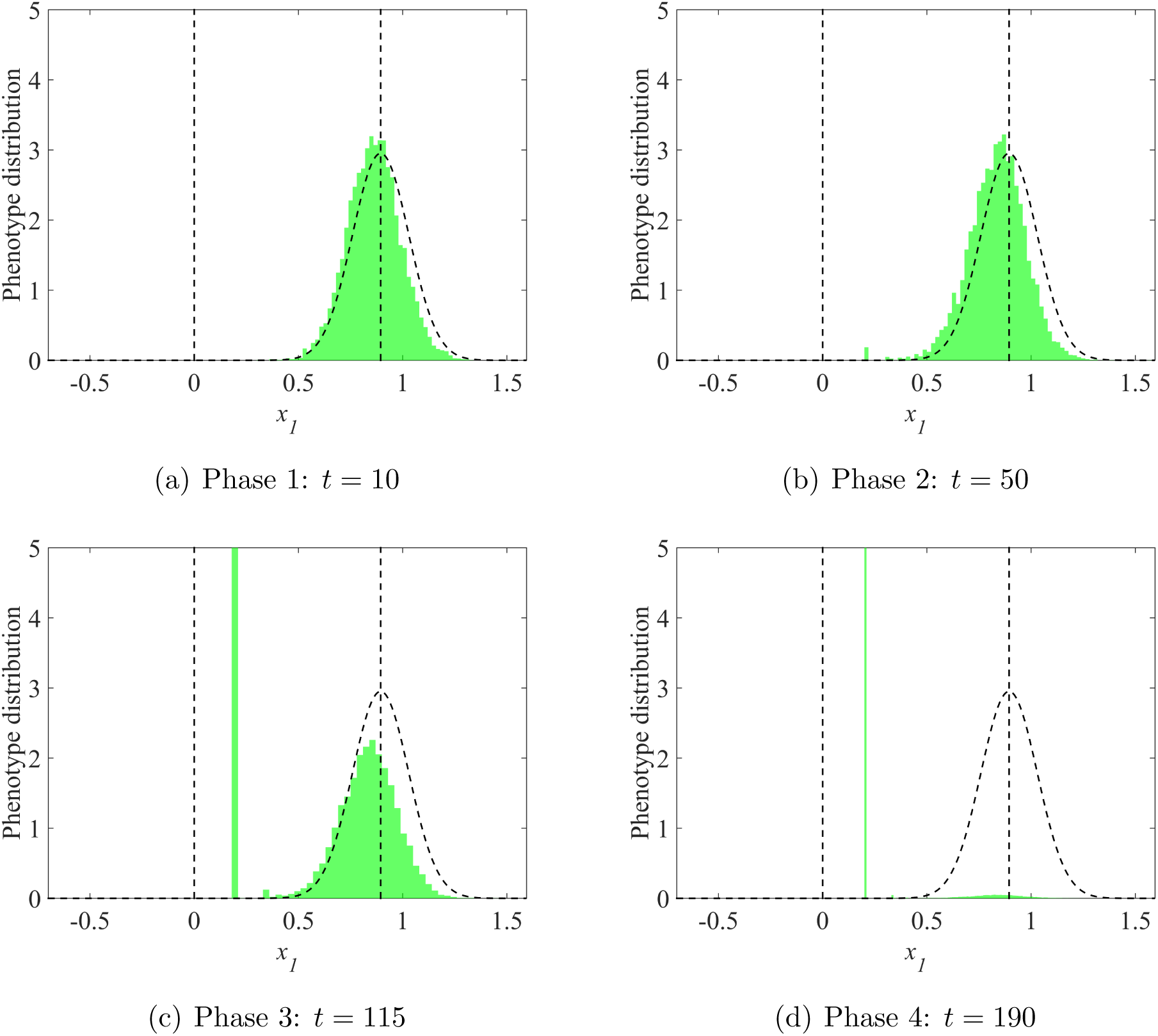
Phenotype distribution in the sink, along the direction *x*_1_, in the absence of mutation. The vertical dotted lines correspond to the sink (*x*_1_ = 0) and source 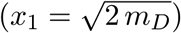 optima. The black dotted curve corresponds to the theoretical distribution of migrant’s phenotypes in the sink (Gaussian distribution, centered at 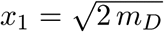, and with variance 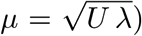). The parameter values are *m*_*D*_ = 0.4, *U* = 0.1, *r*_max_ = 0.1, *λ* = 1*/*300, *n* = 6 and *d* = 10^4^.

