## Supplementary File 1 for "When sinks become sources: adaptive colonization in asexuals": FGM_intermediate_sink.m

```
% Simulation of source-sink dynamics in the presence of an intermediate
% habitat
% version: 16 jan 2019
% Authors:
% Cite: Lavigne, Martin, Anciaux, Papaix, Roques, When sink become sources.

clear all
close all

%%% loads the source population (phenotypes Gs, and parameters: N, U, n,
%%% lambda, mD) created with FGM_source_SI
try
    load('pheno_source')
catch
    display('Launch FGM_source_SI first')
    break
end

%%% other parameters
rmax=0.1; % maximum absolute fitness in the sink
d=1e4; % immigration rate (source->intermediate, source->  sink, intermediate -> sink)
K=N; %  carrying capacity in the intermediate habitat (=N^*)
T=5e3; % maximum number of iterations

%%%  Controls random number generation
rand('state',sum(100*clock));
randn('state',sum(100*clock));

%%% positions of the optima (overall harshness of stress mD=mD(z^*->0)=mD(z^*->z^**)+mD(z^**->0))
% gstar1 : optimum z^* in the source
gstar1=zeros(1,n);
gstar1(1)=-sqrt(2*mD);
% gstar2 : optimum z^{**} in the intermediate sink
gstar2=zeros(1,n);
gstar2(1)=-sqrt(2*mD)/2;
gstar2(2)=-sqrt(2*mD)/2;
% gstar3 : optimum in the final sink
gstar3=zeros(1,n);
% matrix with the 3 optima
GSTAR(1,:)= gstar1;
GSTAR(2,:)= gstar2;
GSTAR(3,:)= gstar3;

%%% initializations

% population sizes
NV=zeros(3,1); NV(1)=K;
% phenotypes in each environment
G=cell(3,1);
G{1}=Gs;
%  estblishment times in each sink (initialization)
t0(2)=T; t0(3)=T;


%%% main loop
t=0; stop=0;
while (stop==0)&&(t<=T)
    t=t+1;
    % number of phenotypes in each environment
    pop_number(1)=K;
    pop_number(2)=size(G{2},1);
    pop_number(3)=size(G{3},1);
    %%% migration step
    % migrants from the Source to intermediate sink
    nSI=poissrnd(d);
    CMSI=[];
    if nSI>0
        indm=randsample(pop_number(1),nSI); % corresponding indices
        Gi=G{1};
        CMSI=Gi(indm,:); % matrix with the migrant phenotypes
    end
    % number of migrants from the Source to final sink
    nSS=poissrnd(d);
    CMSS=[];
    if nSS>0
        indm=randsample(pop_number(1),nSS); % corresponding indices
        Gi=G{1};
        CMSS=Gi(indm,:); % matrix with the migrant phenotypes
    end
    % number of migrants from the intermediate to final sink
    nIS=poissrnd(d*pop_number(2)/K);
    CMIS=[];
    if nIS>0
        indm=randsample(pop_number(2),nIS); % corresponding indices
        Gi=G{2};
        CMIS=Gi(indm,:); % matrix with the migrant phenotypes
        Gi(indm,:)=[]; % migrants are removed from intermediate sink
        G{2}=Gi;
    end
    % migrants are added to the corresponding environments
    G{2}=[G{2}; CMSI];
    G{3}=[G{3}; CMSS ; CMIS];
    
    % pop sizes after migration
    for i=1:3
        pop_number(i) = size(G{i},1);
    end
    
    %%% reproduction/selection step
    % intermediate sink
    if pop_number(2)>0
        i=2;
        Gi=G{i};
        Mi=sum(-(Gi-repmat(GSTAR(i,:),pop_number(i),1)).^2/2,2); % relative fitness
        rMi=rmax+Mi; % absolute fitness
        ndesc=poissrnd(sum(exp(rMi))); % number of offspring
        V=randsample(pop_number(i),min(K,ndesc),true,exp(rMi)); % sample min(K,ndesc) individuals among the parents (with replacement)
        Gi=Gi(V,:); % new phenotypes after reproduction step
        G{2}=Gi;
        pop_number(2) = size(G{2},1);
    end
    % final sink
    if pop_number(3)>0
        i=3;
        Gi=G{i};
        Mi=sum(-(Gi-repmat(GSTAR(i,:),pop_number(i),1)).^2/2,2); % relative fitness
        rMi=rmax+Mi; % absolute fitness
        ndesc=poissrnd(sum(exp(rMi))); % number of offspring
        V=randsample(pop_number(i),min(K,ndesc),true,exp(rMi)); % sample min(K,ndesc) individuals among the parents (with replacement)
        Gi=Gi(V,:); % new phenotypes after reproduction step
        G{3}=Gi;
        pop_number(3) = size(G{3},1);
    end
    
    %%% mutation step
    for i=2:3
        if pop_number(i)>0
            Gi=G{i}; % phenotypes environment i
            J=poissrnd(U,[1 pop_number(i)]);
            if sum(J>0)>1
                dG1=randn(sum(J>0),n);
                % individuals with at least one mutation
                dG2=repmat(sqrt(J(J>0)*lambda)',1,n); % mutational variance is taken into account
                dG3=dG1.*dG2; % effect of mutations : mutational variance and number of mutations
                dG=zeros(pop_number(i),n);
                dG(J>0,:)=dG3; % matrix of size pop_number(i) x n with the mutation effects
                Gi=Gi+dG; % new phenotypes after mutation step
                G{i}=Gi;
            end
        end
    end
    
    %%% computation of t0
    
    for i=2:3
        if (t0(i)==T)&&(pop_number(i)>0)
            Mi=sum(-(G{i}-repmat(GSTAR(i,:),size(G{i},1),1)).^2/2,2); % relative fitness
            rMi=rmax+Mi; % absolute fitness
            t0(i)=t*(mean(rMi)>0)+T*(mean(rMi)<=0); % t0(i) is the first time when the absolute fitness becomes positive in environment i
        end
    end
    
    % stopping criterion
    if t0(3)<T||(t>T+1)
        stop=1;
    end
    
    % histogram with the population sizes
    if mod(t,10)==0
        bar(pop_number)
        title(['t=' num2str(t)])
        set(gca,'xtick',[1  2 3])
        set(gca,'xticklabel',{'source'  'inter' 'sink'})
        ylabel('pop. sizes','FontSize',20)
        pause(0.1)
    end
end

display(['Time needed to invade the intermediate sink: ' num2str(t0(2))])

display(['Time needed to invade the sink: ' num2str(t0(3))])
```
