## Supplementary File 1 for "When sinks become sources: adaptive colonization in asexuals": FGM_source_SI.m

```
% Simulation of a source population, at mutation-selection balance
% version: 16 jan 2019
% Authors:
% Cite: Lavigne, Martin, Anciaux, Papaix, Roques, When sink become sources.

clear all
close all

%%% parameters
N=1e6; % population size
U=0.1; % mutation rate
n=6; % phenotype dimension
lambda=1/300; % mutational variance
mD=0.4; % harshness of stress

%%% number of generations
mu=sqrt(U*lambda); % mutational parameter
T=20/sqrt(mu); % overestimation of the time needed to reach mutation-selection balance

%%%  Controls random number generation (random seed, based on internal
%%%  clock)
rand('state',sum(100*clock));
randn('state',sum(100*clock));

%%% initialization: clonal population, at the phenotype optimum (x^*)
gstar1=zeros(1,n); gstar1(1)=-sqrt(2*mD);  % gstar1: optimum x^* in the source
Xs=repmat(gstar1,N,1); % replicate N times
Gs=Xs; % matrix containing all of the phenotypes in the population

%%% main loop
for t=1:T
    % Darwinian fitness
    Q=sum(-(Gs-Xs).^2/2,2);    M=exp(Q);
    % reproduction and selection
    V=randsample(N,N,true,M); % draw N phenotypes among the population of size N, with probability of being sampled given by M
    Gs=Gs(V,:); % New population, after the reproduction and selection step
    % number of mutations (vector of length N)
    J=poissrnd(U,[1 N]);
    % computation of the mutation effects on phenotypes
    if sum(J>0)>0
        dG1=randn(sum(J>0),n); % mutation effects, without taking into account the number of mutations and the variance
        dG2=repmat(sqrt(J(J>0)*lambda)',1,n); dG3=dG1.*dG2; % the variance and the number of mutations are taken into account
        dG=zeros(N,n); dG(J>0,:)=dG3; Gs=Gs+dG; % phenotypes are modified accordingly
    end
    % plots the distribution of fitness
    plotx=0; %
    if (plotx==1)
        histogram(Q,[-0.2:0.001:0],'Normalization','pdf','Edgecolor','none'); xlim([-0.2 0.05]); ylim([0 40]);
        xlabel('m','FontName','Times New Roman','FontAngle','italic','FontSize',16); set(gca,'FontSize',16,'FontName','Times New Roman');
        pause(0.1)
    end
end

%%% saves the matrix with the phenotypes and the parameters (to be used in FGM_source_sink_SI)
save pheno_source Gs N U n lambda mu mD
```
