## Supplementary File 1 for "When sinks become sources: adaptive colonization in asexuals": FGM_source_sink_SI.m

```
% Simulation of source-sink dynamics
% version: 16 jan 2019
% Authors:
% Cite: Lavigne, Martin, Anciaux, Papaix, Roques, When sink become sources.

clear all
close all

%%% loads the source population (phenotypes Gs, and parameters: N, U, n,
%%% lambda, mD) created with FGM_source_SI
try
    load('pheno_source')
catch
    display('Launch FGM_source_SI first')
    break
end
%%% other parameters
rmax=0.1; % maximum absolute fitness in the sink
d=1e4; % immigration rate

%%%  Controls random number generation (random seed, based on internal
%%%  clock)
rand('state',sum(100*clock));
randn('state',sum(100*clock));

%%% initialization
Nt=0; % population size in the sink
G=[]; % corresponding phenotypes
arret=0;
t=0;

%%% boucle principale
while arret==0
    t=t+1;
    %%% immigration stage
    Nm=poissrnd(d); % number of migrants (Poisson distribution with mean d)
    if Nm>0
        Lm=randi([1,N],Nm,1); % draw Nm individuals in the source population
        Gm=Gs(Lm,:); % genotypes of the migrants
        G=[G ; Gm]; % migrants are included in the sink
        Nt=Nt+Nm; % new population size
    end
    
    if Nt>0
        %%% reproduction
        % Darwinian fitness
        Q=sum(-(G-0).^2/2,2); M=exp(Q); % Q is the Malthusian fitness
        % mean relative fitness
        m(t)=mean(Q);
        % new population size, after reproduction
        Nt1= poissrnd(sum(M*exp(rmax))); % sum of Poisson random variables
        
        %%% adaptation
        % selection
        V=randsample(Nt,Nt1,true,M); % draw Nt1 phenotypes among the population of size Nt, with probability of being sampled given by M
        Nt=Nt1; % updates the population size
        G=G(V,:); % and the phenotypes
        % number of mutations (vector of length Nt)
        J=poissrnd(U,[1 Nt]);
        if sum(J>0)>0
            dG1=randn(sum(J>0),n);  % mutation effects, without taking into account the number of mutations and the variance
            dG2=repmat(sqrt(J(J>0)*lambda)',1,n); dG3=dG1.*dG2;  % the variance and the number of mutations are taken into account
            dG=zeros(Nt,n); dG(J>0,:)=dG3; G=G+dG; % phenotypes are modified accordingly
        end
        NNt(t)=Nt; % saves the population size at time t
    else % if Nt==0
        m(t)=NaN;
        v(t)=NaN;
        NNt(t)=Nt;
    end
    %%% stopping criterion
    if t>5e3
        arret=-1;
    elseif Nt>1.5e6
        arret=-2;
    end
    %%% plot
    plotx=0; %
    if (plotx==1)
        histogram(Q,[-0.5:0.001:0],'Normalization','pdf','Edgecolor','none'); xlim([-0.2 0.05]); ylim([0 40]);
        xlabel('m','FontName','Times New Roman','FontAngle','italic','FontSize',16); set(gca,'FontSize',16,'FontName','Times New Roman');
        pause(0.1)
    end
end
T=t; % final time

%%% plots the trajectory of mean fitness and population size

%%% trajectory of mean fitness
figure 
% rbar obtained by IBM simulation
plot(m+rmax,'c','LineWidth',1.5)
xlabel('t','FontName','Times New Roman','FontAngle','italic','FontSize',20)
ylabel('$\overline{r}(t)$','Interpreter','latex','FontSize',20)
set(gca,'FontSize',16,'FontName','Times New Roman')
hold on
% analytical value of rbar (black curve)
transf=@(z)tanh(mu*z)/mu; f=@(z) exp(mD*mu*transf(z).^2./(1+mu*transf(z))-mD*transf(z)+(rmax-n/2*mu)*z); Den=@(z)integral(f,0,z); mt=[];
for t=1:T
    aux=((f(t)-1)./Den(t))-rmax;
    mt=[mt;aux];
end
plot(mt+rmax,'k-','LineWidth',1.5)
grid


%%% population size
figure
% N obtained by IBM simulation
semilogy(NNt,'color',[1 0.5 0.5],'LineWidth',1.5);
ylim([1e3 2e6])
xlabel('t','FontName','Times New Roman','FontAngle','italic','FontSize',20)
ylabel('$N(t)$','Interpreter','latex','FontSize',20)
set(gca,'FontSize',16,'FontName','Times New Roman')
hold on
% analytical value of N
Nt(1)=d*exp((rmax-mD-mu*n/2)); 
for t=1:length(m)-1
    Nt(t+1)=(Nt(t)+d)/(1-(m(t+1)+rmax));
end
semilogy(Nt,'k-','LineWidth',1.5)
grid
```
